# Phylogenomics of a genus of ‘Great Speciators’ reveals rampant incomplete lineage sorting, gene flow, and mitochondrial capture in island systems

**DOI:** 10.1101/2024.08.28.610082

**Authors:** Jenna M. McCullough, Chad M. Eliason, Shannon Hackett, Corinne E. Myers, Michael J. Andersen

**Author notes:** Corresponding author: Jenna McCullough.

## Abstract

The flora and fauna of island systems, especially those in the Indo-Pacific, are renowned for their high diversification rates and outsized contribution to the development of evolutionary theories. The total diversity of geographic radiations of many Indo-Pacific fauna is often incompletely sampled in phylogenetic studies due to the difficulty in obtaining single island endemic forms across the Pacific and the relatively poor performance of degraded DNA when using museum specimens for inference of evolutionary relationships. New methods for production and analysis of genome-wide datasets sourced from degraded DNA are facilitating insights into the complex evolutionary histories of these influential island faunas. Here, we leverage whole genome resequencing (20X average coverage) and extensive sampling of all taxonomic diversity within *Todiramphus* kingfishers, a rapid radiation of largely island endemic ‘Great Speciators.’ We find that whole genome datasets do not outright resolve the evolutionary relationships of this clade: four types of molecular markers (UCEs, BUSCOs, SNPs, and mtDNA) and tree building methods did not find a single well-supported and concordant species-level topology. We then uncover evidence of widespread incomplete lineage sorting and both ancient and contemporary gene flow and demonstrate how these factors contribute to conflicting evolutionary histories. Our complete taxonomic sampling allowed us to further identify a novel case of mitochondrial capture between two allopatric species, suggesting a potential historical (but since lost) hybrid zone as islands were successively colonized. Taken together, these results highlight how increased genomic and taxon sampling can reveal complex evolutionary patterns in rapid island radiations.

---

Islands play an outsized role in the study of speciation of wild populations (Wallace 1881; MacArthur and Wilson 1967; Whittaker et al. 2017). Indeed, some of the most influential examples of species diversification stem from island systems. These include the adaptive radiations of Darwin’s finches in the Galapagos (Darwin 1859; Grant and Grant 2006; Lamichhaney et al. 2015), Hawaiian silverswords and honeycreepers (Baldwin and Sanderson 1998; Lerner et al. 2011; Landis et al. 2018), and Caribbean *Anolis* lizards (Mahler et al. 2010). Though these examples have provided substantial insights in evolutionary biology, they are unique cases in which results may not be generalizable to other groups of organisms. Conversely, many lineages have evolved equally rapidly on islands without morphological novelty or obvious adaptive consequences—hereafter referred to as geographic radiations (Rundell and Price 2009; Simões et al. 2016). As distinct from adaptive radiations, geographic radiations occur across spatially structured landscapes, such as island archipelagos, without the evolution of key innovations or obvious adaptive changes (Moyle et al. 2009; Lieberman 2012; Moen and Morlon 2014). By studying organisms that occur across island systems, researchers can more readily parse how these forces shape the evolutionary history of clades during bouts of rapid and successive speciation events.

However, the study of rapid geographic radiations across island systems is not without its challenges. For example, neutral processes like genetic drift and mutation are stronger evolutionary forces in small populations such as those found in island systems (Baum et al. 2017). Additionally, rapid and successive speciation events contribute to short internode distances that prove difficult for phylogenetic inferences (Whitfield and Lockhart 2007; Bagley et al. 2020; Roycroft et al. 2020). These events also contribute to incomplete lineage sorting (ILS), which is more frequent when successive speciation events occur at a faster rate than the coalescence of alleles (Degnan and Salter 2005; Maddison and Knowles 2006; Scornavacca and Galtier 2017). These issues are magnified when historical samples are included in analyses (Smith et al. 2020; Ernst et al. 2022; Roycroft et al. 2022), a necessary, yet adverse, byproduct of the gaps in modern genetic sampling. As our ability to collect better genetic data from degraded samples increases, empirical studies are turning to historical samples to include these range-restricted or rarely collected populations to broadly sample the entirety of geographic radiations.

The varied geographic radiations across the Indo-Pacific, encompassing southeast Asia eastward to Polynesia, have contributed to many aspects of ecological and evolutionary theories in wild populations. These include the early foundations of biogeography (Müller 1846; Wallace 1863, 1881; Lydekker 1896), the Theory of Island Biogeography (MacArthur & Wilson, 1967), the ‘Great Speciator’ paradox (Diamond et al., 1976), taxon cycles (Wilson, 1959, 1961), and allopatric speciation (Mayr 1942, 1963). Considered to be the main mechanism of speciation, allopatric speciation invokes geographic isolation as a critical aspect of reproductive isolation and subsequent lineage diversification. Prior to his work on speciation and the Biological Species concept (Dobzhansky 1937; BSC; Mayr 1940, 1942), Ernst Mayr wrote extensively on the phenotypic differences and evolutionary histories of many avian geographic radiations as curator of ornithology at the American Museum of Natural History. He was particularly interested in describing geographic variation based on avian specimens collected during the Whitney South Seas Expedition (Chapman 1935), including honeyeaters (Mayr 1932a), *Rhipidura* fantails (Mayr 1931a), *Pachycephala* whistlers (Mayr 1932b), and kingfishers (Mayr 1931b, 1935, 1941). The extensive phenotypic variation across these geographic radiations was influential in the development of his evolutionary theories (Mayr 1940, 1942, 1963). Mayr’s work on the geographic variation of *Halcyon* kingfishers—now *Todiramphus* (Moyle 2006)—led to his coinage of the ‘superspecies’ concept (Mayr 1931b), or what we now refer to as geographic radiations. The superspecies concept describes a taxonomic ‘grey zone’ in which the varied insular populations that comprise a geographic radiation may be recently diverged, having developed distinctive phenotypic characteristics in isolation; however, they could not be treated as separate species given our inability to assess reproductive isolation between allopatric populations, a requirement of the Biological Species concept (Dobzhansky 1937; Mayr 1942, 1963; Mayr and Diamond 2001).

Due to changing prevailing definitions of species and new advances in molecular systematics, there have been drastic impacts on the taxonomy of geographic radiations and subsequent estimations of their species richness. Alternative views on species boundaries, such as the Generalized Lineage Species Concept (de Quieroz 1998; de Queiroz 1999), led to an increase in recognized species-level diversity within allopatric superspecies complexes. For example, there has been a sea change in the way species limits are applied to insular geographic radiations, due to the resolution of paraphyly or recognition of phylogenetically divergent lineages, including in whistlers (Andersen et al. 2014; Marki et al. 2018), thrushes (Reeve et al. 2023), pittas (Irestedt et al. 2013), Australasian robins (Kearns et al. 2016), monarch flycatchers (Andersen et al. 2015a, 2021; McCullough et al. 2021), white-eyes (Lim et al. 2019), sunbirds (Ó Marcaigh et al. 2022), and kingfishers (Andersen et al. 2013; Sin et al. 2022). Increasingly, our understanding of speciation with gene flow in these radiations has brought a more nuanced approach to species limits (Nosil 2008; Campbell et al. 2018; Gyllenhaal et al. 2020; Mapel et al. 2021).

*Todiramphus* kingfishers represent the most species-rich and widespread genus within its taxonomic order (Aves: Coraciiformes), distributed from the Red Sea to French Polynesia (Fig. 1). They are generalist predators, not obligate plunge-diving piscivores, as their common name suggests. The Collared Kingfisher (*Todiramphus chloris*) is among the most widespread and polytypic avian superspecies complexes in the Indo-Pacific. Andersen et al. (2015b) argued for a revised classification based on a phylogeny that found no less than 10 species embedded within the *T. chloris* species complex, thus rendering *T. chloris* paraphyletic. Moreover, this study found that multiple instances of sympatry and obvious reproductive isolation within this group suggested that species limits within *Todiramphus*, and especially within the *T. chloris* species complex, were inadequately circumscribed.

**Figure 1.**
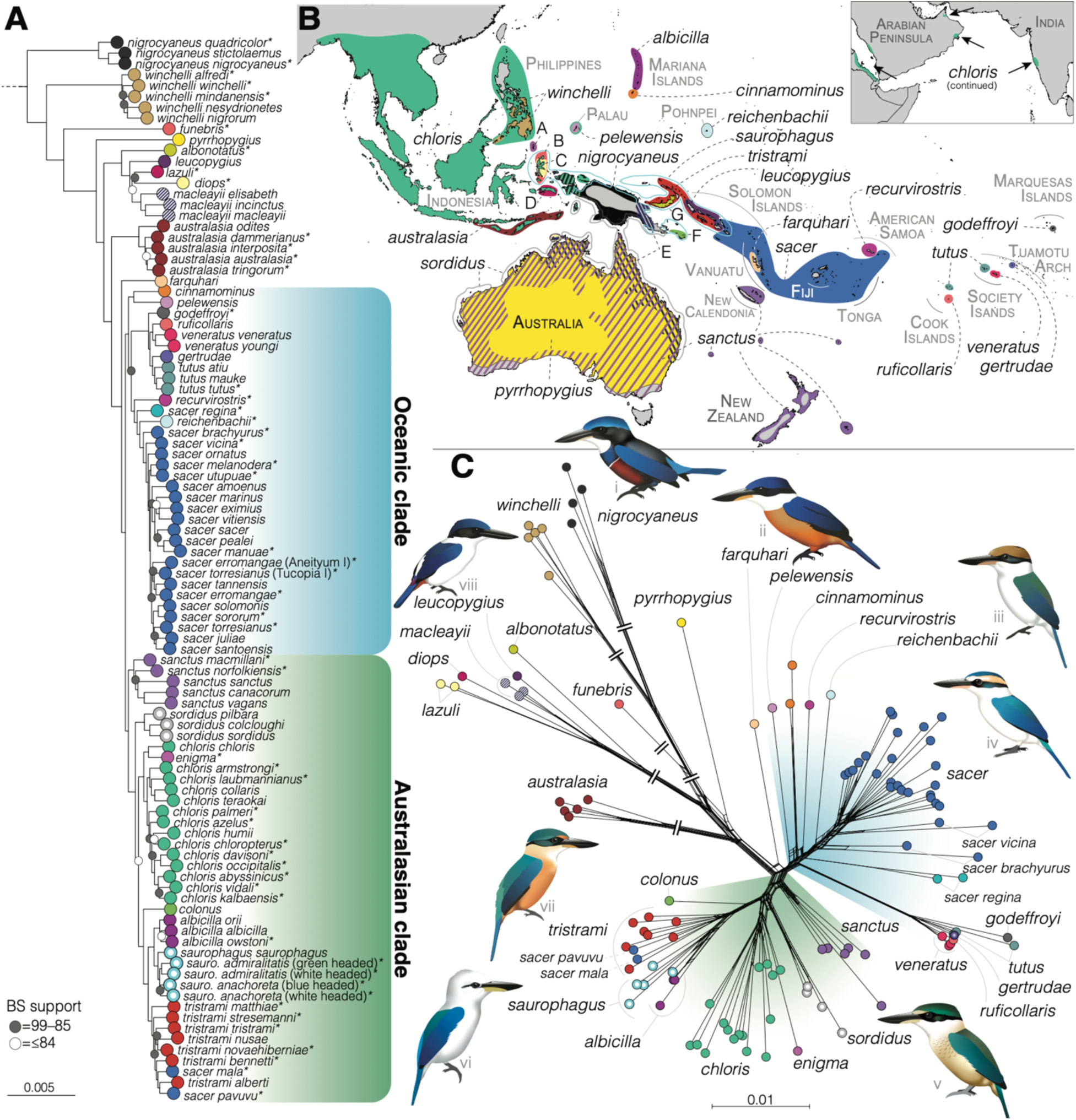
Ranges and evolutionary relationships of all *Todiramphus* kingfishers. A) Partitioned maximum likelihood phylogeny of 4,942 UCE loci from the 90% complete matrix. Circles on nodes indicate support values less than 100 bootstrap support. B). Ranges of 29 species-level groups with diagonal lines showing sympatry and colors corresponding to tips. Letters corresponding to species ranges are as follows: A, *T. enigma;* B, *T. funebris;* C, *T. diops;* D, *T. lazuli*; E, *T. macleayii*, F, *T. colonus*, G, *T. albonotatus*. See Figure S1 with these ranges directly labeled with taxon names. C) Unrooted phylogenetic network based on pairwise divergence among all samples, produced from a complete set of 85,165 SNPs. Note that some branch lengths have been artificially shortened, denoted by double slashes. See Figure S17 for subspecific labels. Illustrations created by Jenna McCullough and names are as follows: i, *T. nigrocyaneus*; ii, *T. farquhari*; iii, *T. reichenbachii;* iv, *T. sacer*; v, *T. sanctus*; vi, *T. saurophagus*; vii, *T. tristrami*; and viii, *T. leucopygius*.

Subsequent analyses highlighted the rapid diversification rate of *Todiramphus* kingfishers (Andersen et al. 2018; McCullough et al. 2019), rivaling the textbook geographic radiation of *Zosterops* white-eyes (Moyle et al. 2009; Lim et al. 2019). Unsurprisingly, the phylogenetic history of *Todiramphus* kingfishers has proven recalcitrant, likely owing to their high speciation rate and relative limitations of prior Sanger and reduced-representation sequencing datasets (Andersen et al. 2015b, 2018; McCullough et al. 2019). More focused work employing RADseq data, particularly assessing a major clade within the genus once considered a part of the *T. chloris* complex, has highlighted a case of mitochondrial discordance within one clade of *Todiramphus* comprising the migratory Sacred Kingfisher (*T. sanctus*), Islet Kingfisher (*T. colonus*), and Torresian Kingfisher (*T. sordidus*; DeRaad et al. 2023). However, this study found negligible levels of gene flow between these species and instead showed patterns consistent with high rates of incomplete lineage sorting.

Because of the relatively rapid timing and phylogenetic diversity in the group, *Todiramphus* kingfishers provide an opportunity to investigate how and when rapid radiations occur across the Indo-Pacific. *Todiramphus* kingfishers originated in Sahul (Australia and New Guinea; Andersen et al. 2018), having diverged from its sister taxon *Syma* approximately 10.3 Ma (McCullough et al. 2019). Since then, they dispersed eastward to French Polynesia and westward across Southeast Asia to western India and as far as the Arabian Peninsula (Fig. 1). The genus exhibits a high degree of secondary sympatry in island systems. For example, five *Todiramphus* species can be found on Halmahera Island, Indonesia (17,780 km^2^; Woodall 2001), and three species can be found on the much smaller Tetepare Island, Solomon Islands (118 km^2^; McCullough et al. 2023). These kingfishers show marked geographic diversity, with noticeable differences in plumages and body size within the same archipelago (e.g., Cinnamon-banded Kingfisher *T. australasia* in the Lesser Sundas) or continental landmasses (Blue-black Kingfisher *T. nigrocyaneus* in New Guinea). Of the 29 currently recognized extant species of *Todiramphus* (Clements et al.2023; Gill et al. 2024), 28 occur on islands and seven are single-island endemics (Fry et al. 1992; Woodall 2001). At the subspecific level, for which there are 93 named taxa, 70 occur only on islands and 28 are restricted to single islands (Fig. S1). Indeed, they occur on nearly every island from Southeast Asia, through Wallacea, Melanesia, Micronesia, and Polynesia—they truly are a quintessential geographic radiation.

Various studies have addressed species-level relationships at the family and ordinal levels (Moyle 2006; Andersen et al. 2018; McCullough et al. 2019) or have addressed specific clades (Andersen et al. 2015b; O’Connell et al. 2019; DeRaad et al. 2023). Yet to date, only half of the total taxonomic diversity within *Todiramphus* has been included in a phylogenetic analysis due to the lack of accessible modern tissue samples. Owing to these sampling gaps, the taxonomy and evolutionary history of *Todiramphus* kingfishers remains unresolved. In this study, we employed whole-genome resequencing of all 93 extant *Todiramphus* taxa to produce the first fully sampled phylogeny of one of Mayr’s classic examples of a geographic radiation. From whole genome resequencing data of 127 samples, we extracted four types of molecular data for phylogenetic inferences: ultraconserved elements (UCEs), single nucleotide polymorphisms (SNPs), Benchmarking Universal Single-Copy Orthologs (BUSCOs), and mitochondrial genomes (mtDNA). In doing so, we provide the most detailed study to date on the evolution of one of the most widespread of the ‘Great Speciator’ genera. Our results reveal how incomplete lineage sorting and gene flow during a rapid radiation have contributed to phylogenetic uncertainty as well as another case of mitochondrial discordance within this complex. Together, our study highlights the complicated evolutionary history of one of the Indo-pacific’s greatest avian radiations.

## METHODS

### Sampling and laboratory methods

We sampled at least one individual for all extant and named *Todiramphus* taxa (Table S1). Because the number of taxa considered for each species varies widely, our species-level sampling ranged from a single sample representing the monotypic Somber Kingfisher (*T. funebris*) to 22 samples representing all the taxa within the widespread Pacific Kingfisher (*T. sacer*). We also sampled two populations of *T. sacer* that Mayr considered as possible separate taxa but refrained from naming both as separate subspecies, from Tucopia Island, Solomon Islands and Aneityum Island, Vanuatu (Mayr 1931b). Moreover, we sampled multiple individuals for two range-restricted Beach Kingfisher taxa that exhibit various head-color morphotypes (*T. saurophagus admiralitis* and *T. s. anachoreta* which have green/blue or white color on their heads). The only *Todiramphus* species we did not sample was the extinct Mangareva Kingfisher (*T. gambieri*), whose species is represented by a single specimen collected from Mangareva Island in 1838 (Thibault and Cibois 2017). Many *Todiramphus* taxa are endemic to a few or single islands across the Indo-Pacific and are thus rarely collected, so we relied heavily on clippings of toepads from museum study skins (n = 76, 59% of total sampling). All other samples were either sourced from blood (n = 4) or ethanol- or frozen-preserved tissue samples (n = 47) loaned to us from natural history collections. Our sampling scheme spans nearly one and a half centuries of collecting effort by countless collectors (1887–2019, toepads span 1887–1987). In total, we had 127 samples representing all 92 extant diversity within the genus; because of differing levels of sample quality, not all samples are present in each generated dataset (Table S1). We chose four representatives from each of the three kingfisher subfamilies as outgroups, sourced from higher-level work on genomic evolution across the family (Eliason et al. 2023). Because DNA sourced from museum specimens is known to have higher rates of degradation than high quality tissues, we treated toepad-sourced samples differently than tissue-sourced samples at all stages of wet lab work (see Appendix 2 for more details). We prepared libraries with Kapa Biosystems Hyper Prep Kit and the iTruStub dual-indexing system ((baddna.org) using established protocols at the University of New Mexico. We pooled toepad- and tissue-sourced libraries separately (10–15 toepad libraries per lane and 15–16 tissue libraries per lane) and sequenced them across 12 S4 PE-150 lanes of an Illumina NovaSeq 6000 at the Oklahoma Medical Research Foundation.

### Bioinformatic processing

We trimmed demultiplexed raw reads with trimmomatic v.0.39 (Bolger et al. 2014) and aligned cleaned reads to a Collared Kingfisher (*T. chloris collaris*) reference genome (Eliason et al. 2022) using BWA v.0.7.17 (Li and Durbin 2009) and GNU Parallel (Tange 2021). We used MarkDuplicatesSpark, implemented in GATK v.4.2.6.1 (McKenna et al. 2010; Van der Auwera and O’Connor 2020), to tag and remove duplicate reads. We assessed individual genome summary statistics (i.e., CollectAlignmentSummaryMetrics, CollectInsertSizeMetrics, and depth) with Picard v.2.26.11 (Broad Institute 2019) and samtools v.1.15 (Danecek et al. 2021). We used GATK v.3.8.1 to identify and realign indels (RealignerTargetCreator, IndelRealigner) to produce final BAM files for extraction of different types of molecular markers for downstream phylogenetic analyses.

### Extraction and analysis of Ultraconserved Elements (UCEs)

We used the Phyluce v.1.7.0 (Faircloth 2016); described in full at https://github.com/faircloth-lab/phyluce) Python package to harvest thousands of UCE loci in silico (specifically following the phyluce Tutorial III: Harvesting UCE Loci From genomes). We used samtools to convert the final BAM file to fasta and then to 2.bit format using fatoTwoBit (Kuhn et al. 2013). Next, we aligned the tetrapods UCE 5k probe set to each genome and extracted loci with 500 bp flanking regions with “phyluce_probe_slice _sequence_from_genomes”. We then followed the typical bioinformatic steps implemented in the phyluce pipeline starting from “phyluce_assembly_get_match_counts.” We aligned each locus with MAFFT (Katoh and Standley 2013) without initially trimming the ends of UCE loci, instead trimming using TrimAL v.1.4.rev15 (Capella-Gutiérrez et al. 2009) with the “-automated1” flag and degenerated_DNA matrix (available at https://github.com/inab/trimal/blob/trimAl/dataset/matrix.Degenerated_DNA) to allow for ambiguous bases.

During preliminary phylogenetic exploration, we determined that many of the toepad-sourced samples had exceptionally long branches (see below for phylogenetic analyses and Table S1 for these samples). Biologically unrealistic long branches such as these can contribute to long-branch attraction and therefore incorrect phylogenetic relationships (Felsenstein 1978); this issue has been shown as a byproduct of poor alignment trimming contributing to “dirty ends” of UCE loci sourced from historical samples, including those sourced from avian toepads (McCormack et al. 2015; Smith et al. 2020; Salter et al. 2022). We removed these artefactual “dirty ends” following the bioinformatic pipeline by Smith et al. (2020; see Appendix 2 for further information). Our final UCE dataset was a 90% complete matrix comprising multiple individuals for some taxa that had at least 105 of 117 samples present at each UCE locus (Table S3).

We used multispecies coalescent tree (MSC) and concatenated maximum likelihood (ML) methods to analyze UCE datasets. Because gene tree histories could conflict with the overall species tree, we used two species tree methods that account for gene tree heterogeneity (Kubatko and Knowles 2023). First, we used SVDquartets (Chifman and Kubatko 2014) implemented in Paup* 4.0a168 (Swofford 2003); this quartet-based method uses a concatenated alignment and site pattern frequencies of splits to estimate a species tree and has been shown to perform well for large multilocus datasets (Wascher and Kubatko 2021). To decrease computation time in Paup*, we split up the 117-sample, 90% complete matrix into two, overlapping subsets of taxa (each with 69 samples). For each subset, we analyzed all possible quartets and assessed support with 100 BS. Next, we used the summary-statistic method ASTRAL-III v.1.15.2.4 (Zhang et al. 2018; Rabiee et al. 2019) to estimate the species tree. This summary-statistic MSC method is sensitive to fragmentary data and depends on high quality gene trees (Mirarab 2023). Because toepad-sourced samples often have a higher degree of missing data within individual alignments, we used a series of filtering steps to produce a final set of high quality gene trees. To do so, we 1) filtered the 90% complete matrix to drop samples from individual alignments if they comprised more than 25% of ambiguous nucleotides (N) and 70% of gaps; 2) filtered our gene trees based on different thresholds of parsimony informative (PI) sites (>50 PI sites and 1,583 gene trees; >100 PI sites and 125 gene trees; see Appendix 2 for more details). For ASTRAL, we increased the number of searches (-r) and subsampling (-s) from the default of 4 to 8. To estimate a concatenated ML phylogeny of the 90% complete matrix, we used RaxML-NG v.1.2.0 (Kozlov et al. 2019), assuming a general time-reversible model of nucleotide substitution and gamma-distributed rates among all sites (GTR+G) and evaluated support with the autoMRE function (set to 1000 BS). Finally, we inferred a partitioned maximum likelihood phylogeny with IQtree 2.1.4 (Chernomor et al. 2016; Minh et al. 2020b), implementing modelfinder and assessing support with 1,000 replicates of Ultrafast bootstraps (Hoang et al. 2018).

### Processing and analysis of single-copy orthologous loci (BUSCOs)

Because of their conserved nature in coding regions of the genome, single-copy orthologous genes (Benchmarking Universal Single-Copy Orthologs [BUSCOs; Simão et al. 2015) have been shown to be informative molecular markers at both deep and shallow timescales (Waterhouse et al. 2018; Timilsena et al. 2022; Alaei Kakhki et al. 2023). Prior to extracting thousands of BUSCO loci for phylogenetic inference, we masked low coverage sites from the final BAM files. Generally, we calculated read depths with mosdepth 0.3.3 (Pedersen and Quinlan 2017), generated consensus sequences with bcftools 1.19 (Li 2011; Danecek and McCarthy 2017) implemented in a custom function in R 4.3.2 (R Development Core Team 2013), and used bedtools v2.31.1 (Quinlan and Hall 2010) to mask sites with less than five reads. We extracted thousands of BUSCO loci from the genomes using BUSCO 5.4.6 (Manni et al. 2021a, 2021b). During initial phylogenomic analysis, it became clear that extracting these data from the lower coverage toepad genomes yielded problematic alignments that contributed to an artificial pattern in which these low coverage toepad genomes grouped together rather than within the clades that matched those inferred with SNP and UCE datastets (Fig. S3). Because all of these data are aligned to the same reference, we instead ran BUSCO on the high quality *T. chloris collaris* reference genome and used BED coordinates of 8,337 BUSCO loci to extract these same regions from all of our samples. To do so, we converted the metaeuk output for our reference sample to gff3 coordinate format using the python script MetaeukToGff.py (https://github.com/Andy-B-123/MetaeukToGff3). With GFFread v0.12.8 (Pertea and Pertea 2020), we extracted individual genes in amino acid (AA) format from all samples. We created locus-specific files with faSomeRecords (Kuhn et al. 2013) and removed short duplicates within loci with keep-longest.py (https://gist.github.com/mkweskin/8869358). Because all loci—regardless of whether adequate coverage of sequence data is present—are extracted based on this approach, we filtered out loci that were entirely composed of missing data or invariant sites as well as those that did not have PI sites to produce a matrix of 8,012 AA loci for 109 samples.

To determine if protein-coding loci inferred similar phylogenetic histories to the SNP and UCE datasets, we also inferred phylogenies for the BUSCO loci with MSC and ML methods. Similar to how we treated UCE data, we performed additional steps to filter BUSCO loci for MSC analyses in order to produce high quality gene trees as input into ASTRAL. Though every sample in our dataset was present at each of these loci regardless of presence of data, many toepad samples comprised 50% or more of ambiguous sites. We dropped these individual samples from BUSCO alignments with the same python script noted in the UCE processing section by the ambiguous character from “N” to “X”. In a similar fashion as the UCE datasets, we filtered BUSCO loci by the number of PI sites. With a threshold of 20 PI AA sites (4,177 gene trees), a similar artefactual pattern of toepad-sourced samples cluster was apparent (Fig. S4) that disappeared with a threshold of 100 PI AA sites (470 gene trees). We increased the number of searches (-r) and subsampling (-s) to 8. We inferred a maximum likelihood phylogeny for the concatenated matrix of 8,012 loci with IQtree 2.1.4 and assessed support with 1,000 ultrafast bootstraps. During model selection, we confined substitution models to the Q.bird matrix and did not allow for partition merging (Chernomor et al. 2016; Kalyaanamoorthy et al. 2017; Minh et al. 2021).

### Processing and analysis of Single Nucleotide Polymorphisms (SNPs)

We performed variant discovery and genotyping on realigned BAM files with UnifiedGenotyper in GATK 3 using a scatter-gather approach. Recent work has shown that the results of this program are comparable to HaplotypeCaller in GATK 4 and ANGSD (Korneliussen et al. 2014) for non-model organisms (Aguillon et al. 2021). With VCFtools, we initially filtered SNPs that were successfully genotyped in at least 90% of samples, had a minimum quality score of 30, minimum depth of 3, and a minor allele count of 3. Because toepad-sourced samples had significantly higher missing data, we removed samples that were missing at 55% or more sites in this initial dataset. This only affected poorly sequenced toepad-sourced samples in which we had several representatives per taxon. One sample of *T. n. nigrocyaneus* (AMNH 301866) fell marginally below this threshold, but since it had less missing data than our other *T. n. nigrocyaneus* sample, thus we still included it in the final VCF file (see Table S1). After dropping these six toepad samples, we then produced a SNP dataset in which all samples were successfully genotyped at all sites, as well as only keeping bi-allelic SNPs every 5,000 bp. Our final dataset comprised 85,156 SNPs from 119 samples representing all *Todiramphus* taxa.

We implemented three different ML, MSC, and distance-based network methods for tree building using SNP data. First, we converted the VCF file to nexus format with nucleotide data using the ruby script convert_vcf_to_nexus.rb (available on https://github.com/mmatschiner/tutorials/blob/master/species_tree_inference_with_snp_data/src/convert_vcf_to_nexus.rb). We used this alignment for ML inference in IQtree, correcting for ascertainment bias (+ASC) to limit branch length overestimations and assessed support with 1,000 BS replicates. Second, we analyzed the SNP alignment using all possible quartets with SVQquartets and assessed support with 1,000 BS replicates. Third, we generated a phylogenetic network to visualize the genetic distances between all samples. We used R package StAMPP v1.6.3 (Pembleton et al. 2013; R Development Core Team 2013) to produce a pairwise genetic distance (Nei’s D; Nei 1972) matrix as input into SplitsTree4 v 4.18.13 (Bryant and Moulton 2004; Huson and Bryant 2006) to produce a Neighbor-Net phylogram.

### Extraction and analysis of Mitochondrial genomes

We generated mitogenomes by mapping the trimmomatic cleaned reads to a reference mitogenome of *T. sanctus vagans* (GenBank: NC011712, (Pratt et al. 2009) using the singularity version of Mitofinder V1.4.1 (Allio et al. 2020). We checked these alignments by eye for incorrect internal stop codons and dropped spurious samples from gene alignments if initial phylogenetic analyses inferred them as an outgroup to *Todiramphus* (suggesting a incorrectly mapped NUMT sequence rather than true mitochondrial DNA; (Nacer and Raposo do Amaral 2017). We used IQtree v2.2.2.6 to infer a maximum likelihood phylogeny of the 12 mtDNA genes and assessed support with 1,000 BS replicates.

### Tests for incomplete lineage sorting and gene flow

To test for incomplete lineage sorting, we calculated gene and site concordance factors for the BUSCO and 90% complete UCE matrix trees inferred with IQtree. Gene (gCFs) and site concordance factors (sCFs) are two additional measures for assessing topological support by revealing the level of gene tree species tree discordance in phylogenomic datasets to assess the extent of ILS. We pared down each of these datasets to a subset of 34 tips that encompassed higher-level relationships across *Todiramphus*. To calculate gCFs, we chose to use the more informative gene trees that we produced for ASTRAL in order to limit noise introduced by thousands of low-signal gene trees. We calculated cGFs from UCE-based gene trees with >50 PIsites (1,583 gene trees) and BUSCO-based gene trees with >20 PI AA sites (4,155 gene trees) with the superfluous tips not included in our 34-tip subset dropped ad hoc. We calculated sCFs, using the maximum likelihood implementation with --scfl flag (Minh et al. 2020a; Mo et al. 2023), from the pared down concatenated alignments.

We estimated the phylogenetic network for the UCE gene tree dataset (1,583 gene trees) with Species Networks applying Quartets (SNaQ; Solís-Lemus and Ané 2016) implemented in Julia (Bezanson et al. 2017). SNaQ is a maximum pseudolikelihood approach to infer phylogenetic networks while simultaneously accounting for ILS and gene flow with a coalescent model based on quartet concordance factors (Baum 2007). We counted quartets and calculated concordance factors with the ‘countquartetsintrees’ function in PhyloNetworks (Solís-Lemus et al. 2017). After removing our outgroup, the Blue-caped Kingfisher (*Actenoides hombroni*), from our gene trees, our 33-tip dataset comprised 40,920 unique quartets which proved computationally intractable to analyze together. Instead, we produced three subsets of 3,000 non-overlapping randomly selected quartets as input into SNaQ, using *h_m_*= 0–4 with 10 runs each. For each subset, we kept adding edges until the likelihood scores no longer improved and chose the tree with the best overall likelihood (though we report the best tree for each subset).

Mitochondrial data revealed discordant topologies for some *T. sacer* taxa and *T. colonus* and *T. sordidus* (discussed in detail in the Results section); to specifically test for gene flow between non-sister taxa, we used Dsuite v0.5 (Malinsky et al. 2021) to perform four-taxon ABBA-BABA tests. We used our 119-sample filtered SNP dataset, in which each sample was present at a given site and each SNP was at least 5,000 bp apart (85,156 SNPs) as input and used the *Dtrios* module for block Jackknifing of 20 SNP windows across the genome (4,256 SNPs per window) to calculate D and *f_4_*-ratio statistics (Durand et al. 2011; Patterson et al. 2012) for every possible quartet topology. We analyzed two different datasets: 1) a species-level dataset in which all subspecific taxa were lumped into their respective species’ tip, except for *T. sacer regina* (as it was not monophyletic with the rest of *T. sacer*; see Results) and 2) the same as #1 but pared down for fewer species and the treatment of five *T. sacer* taxa that were recovered within the mitochondrial *T. sanctus* clade as separate tips: *T. sacer vicina* and four microendemics of Temotu Province, Solomon Islands, *T s. regina, vicina, ornatus, utupuae,* and *brachyurus*. The remaining *T. sacer* taxa were grouped within a “polynesian” and “melanesian” tip based on the two clades recovered in the UCE IQtree topology (Fig. 1). We incorporated a false discovery rate (FDR)-adjusted statistical significance threshold (Benjamini and Hochberg 1995), which is recommended by the developer when running more than one trio (Malinsky et al. 2021). Our FDR-adjusted *p*-threshold was 0.0000137 (3,654 pairwise tests for 29 tips in both datasets).

## RESULTS

The average coverage of assembled genomes was 20X with differences between toepad- (n = 66, average depth 10X with range of 4.6–27.9X) and tissue-sourced genomes (n = 47, average depth 34.8X, 11.9–98X; Table S1, Fig S5). Tissue-sourced genomes were relatively complete, with an average 96.2% (standard deviation ±0.72%) of the 8,338 single-copy BUSCO loci (Table S4, Figs. S6–7). Toepad-sourced genomes, on the other hand, averaged 55.9 ± 33.7% single-copy BUSCO loci; though this was higher for those with more than 10X average coverage (88 ± 14.9%). Average depth for toepad-sourced genomes did not negatively scale with collection date; perhaps not surprisingly, the relative data quality from the datasets extracted from these genomes did scale with average depth (Fig. S7). Ten of the 127 genomes produced for this project, all sourced from toepad material, were not high enough in coverage to produce adequate data for inclusion in each of the four genetic datasets (we highlight in which datasets samples were present in Table S1).

We extracted several datasets with varying numbers of samples for UCE, BUSCO, SNP, and mtDNA data (Table S3). Our 90% complete UCE matrix with 117 samples comprised 4,942 loci and 4,531,264 bp. Our BUSCO datasets, differing based on thresholds of within-locus PI AA sites, comprised 8,012 (>20 PI sites per locus threshold, total length 5,102,241 AA sites) with 108 samples. Our 100% complete SNP dataset, with one SNP per 5 kb, comprised 85,156 SNPs for 119 samples. We recovered complete or partial mitochondrial genomes for 111 samples.

### Topological conflict between nuclear data types and tree-building methods

The backbone of the *Todiramphus* tree is characterized by many short internodes, suggesting rapid and successive speciation events early in the evolution of this group. We found phylogenetic conflict between datasets and methods due, in part, to these short internodes. When just considering species-level relationships, we did not find a single well-resolved topology that matched across any two datasets or tree-building methods (Fig. 2). The first three main branches of the tree rarely varied across analyses, with Blue-black Kingfisher (*T. nigrocyaneus*), Rufous-lored Kingfisher (*T. winchelli*), and Sombre Kingfisher (*T. funebris*), respectively. Only in one analysis did we find *T. winchelli* fully supported as the first branch of the genus instead of *T. nigrocyaneus* (UCE SVDquartets tree; Fig. S8). We found conflicting phylogenetic signals for the next major branch (Fig. 2), as we either recovered: 1) the Australian endemic Rufous-backed Kingfisher (*T. pyrrhopygius*; Figs. S9–10), 2) a small clade of four species as the next branch (referred to as the “*leucopygius* clade”; Figs. 1, S11–18), or 3) *T. pyrrhopygius* sister to the *leucopygius* clade (Fig. S8). Gene and site concordance factors for the UCE and BUSCO IQtree-inferred topologies revealed that both datatypes strongly favored conflicting topologies: 76% of 1,583 best UCE gene trees and 59% of concatenated 90% complete UCE alignment (4,531,264 bp in length) favored the earlier branching *T. pyrrhopygius* topology whereas a similar percentage of the significantly larger BUSCO dataset (4,155 gene trees and 5,102,241 AA sites) favored the *leucopygius* clade branching prior to *T. pyrrhopygius* (Fig. 3). The Lesser Sundas endemic species, Cinnamon-banded Kingfisher (*T. australasia*) was always supported as sister to the clade comprising the bulk of diversity within the genus: the Vanuatu Kingfisher (*T. farquhari*) and all other species. We consistently found support for two major clades within *Todiramphus*: 1) an entirely oceanic island clade and 2) an Australasian clade (Fig. 1). We hereafter refer to these as the “Oceanic clade” and the “Australasian clade” even though some taxa within the Australasian clade also inhabit oceanic islands.

**Figure 2.**
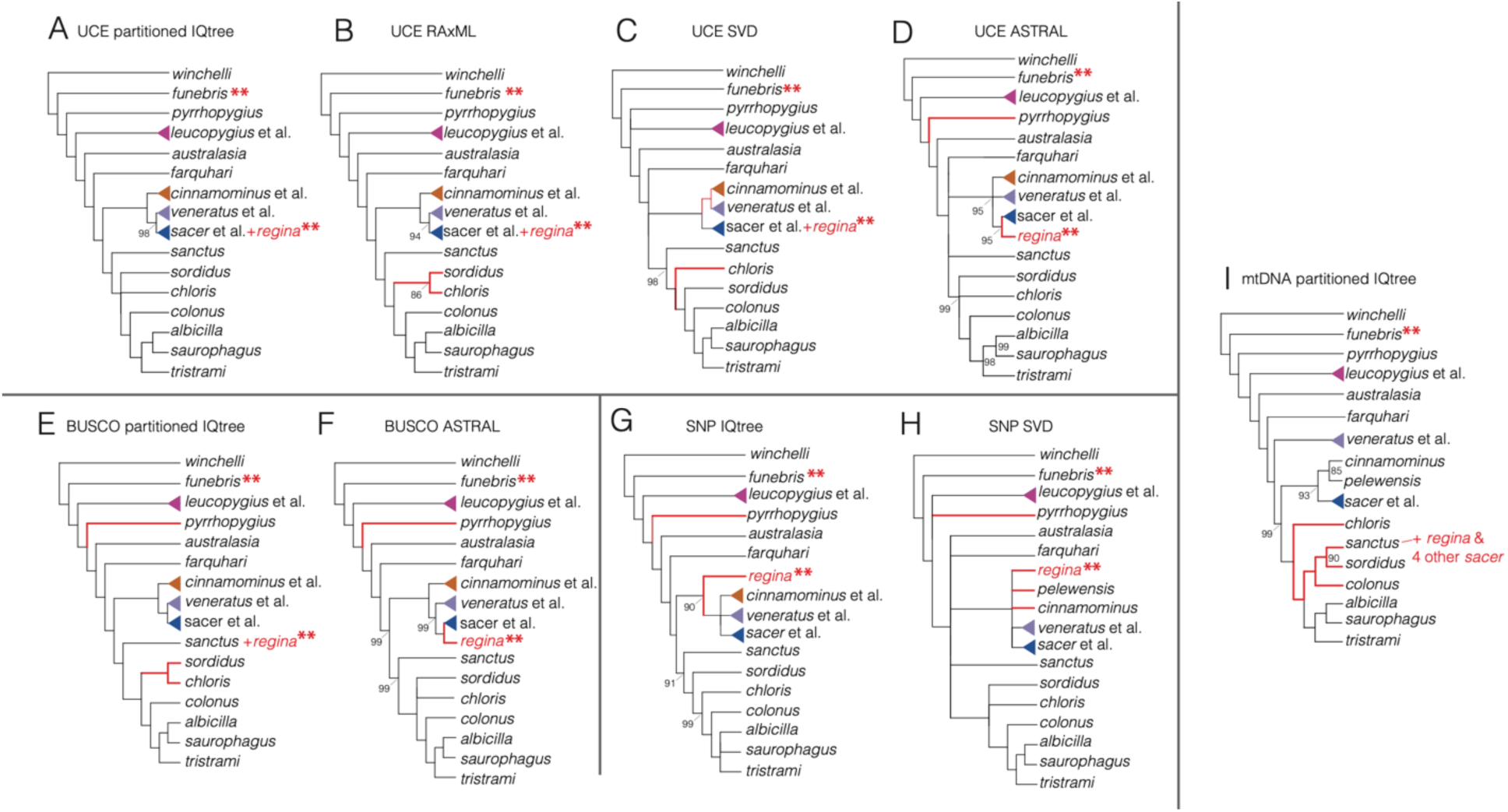
Comparisons of trees from UCE, BUSCO, SNP, and mtDNA datasets with red branches highlighting differences from the UCE maximum likelihood tree. Double asterisks (**) indicate samples sourced from toepad samples. We collapsed nodes with <85 bootstrap support and note those with 99–85 BS. We grouped lineages that were consistently inferred as a clade for simplicity: *T. sacer* et al. includes *T. reichenbachii* and *T. recurvirositris*. Note that *T. pelewensis* and *T. cinnamominus* are considered a single tip (“*cinnamominus* et al..”) except in Panels H and I.

**Figure 3.**
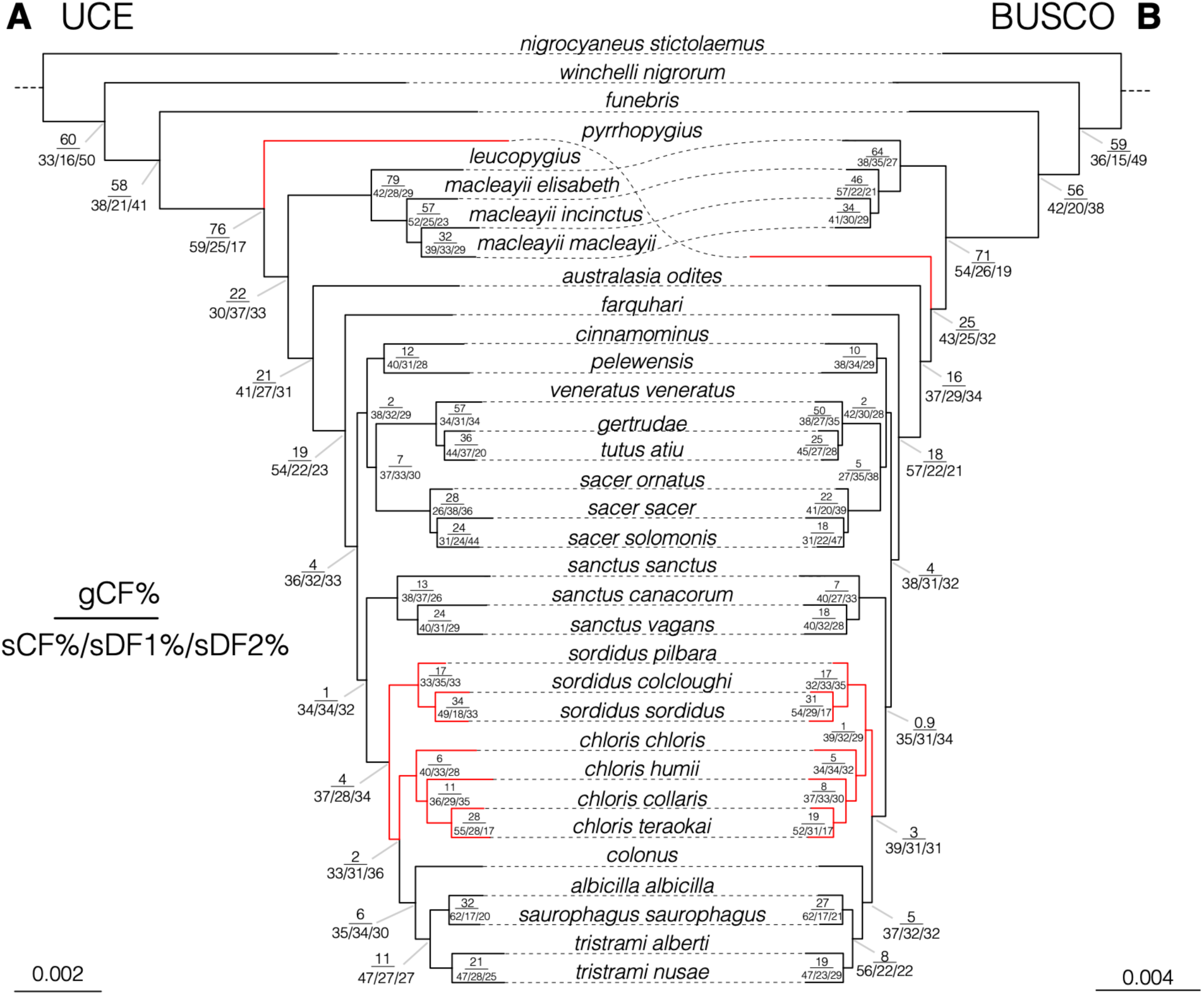
Site (sCF) and gene (gCF) concordance factors of UCE and BUSCO topologies show a high degree of ILS present in the evolutionary history of *Todiramphus* kingfishers. A) Partitioned 90% complete UCE matrix (4,942 loci, total 4,531,264 bp) inferred in IQtree with gCFs calculated from gene trees of loci with at least 50 parsimony informative sites (PIs; 1,583 gene trees). B) Partitioned BUSCO topology (8,012 loci, total 5,102,241 amino acid sites) inferred in IQtree with gCFs calculated from gene trees of loci with >50 PI amino acid sites (4,155 gene trees).

Within the Oceanic clade, we found support for three subclades: 1) a subclade of two microendemics in Micronesia, the Palau Kingfisher (*T. pelewensis*) and the extinct-in-the-wild Guam Kingfisher (*T. cinnamominus*), 2) a subclade of eastern Polynesian species (referred to hereafter as such, comprising *T. veneratus*, *T. gertrudae, T. tutus* etc.), and 3) a subclade comprising Flat-billed Kingfisher (*T. recurvirostris*), Pohnpei Kingfisher (*T. reichenbachii*), and *T. sacer* (i.e., the “*sacer* clade”) sister to the Polynesian subclade. The relationships between these subclades were largely consistent, even though sCF of the relationship between the east Polynesian clade and the *sacer* clade showed that alternative topologies were favored at nearly the same proportion as the inferred topologies for both BUSCO and UCE datasets (Fig. 3). However, most of the discordance between analyses and data types were mostly restricted to whether *T. recurvirostris* and *T. sacer regina* diverged earlier than *T. reichenbachii* and crown *T. sacer* (supported by UCE data; Figs. S8–11, 15) or not (supported by SNP and BUSCO data; Figs. S13–14, 16). Whereas the placement of *T. sacer regina* (Wallis and Futuna), represented by two toepad-sourced genomes (6.8 and 14.5 average coverage), varied between different clades across analyses, we never inferred a topology in which *T. sacer regina* was within crown *T. sacer*. UCE ML analyses instead found *T. s. regina* sandwiched between two other microendemic species in a ladderized branching pattern: *T. recurvirostris* (Samoa), *T. s. regina*, and *T. reichenbachii* (Pohnpei I.; Figs. S1, S9–10). With some support, we also found *T. s. regina*: 1) sister to *T. recurvirostris* (Fig. S8), 2) branching prior to *T. recurvirostris* (Fig. S15), 3) as the first branch of the Oceanic clade (Fig. S14), or 4) as the first branch within the *T. sanctus* complex (Fig. S16). The unrooted phylogenetic network placed *T. sacer regina* as similarly distant from *T. sacer* as *T. recurvirostris* (Figs. 1C; S17).

Within the Australasian clade, eight species formed four subclades: three species complexes (Sacred Kingfisher *T. sanctus,* Torresian Kingfisher *T. sordidus*, and Collared Kingfisher *T. chloris*) and a clade comprising the largely oceanic island species (Islet Kingfisher *T. colonus*, Mariana Kingfisher *T. albicilla*, Beach Kingfisher *T. saurophagus*, and Melanesian Kingfisher *T. tristrami*). *Todiramphus sanctus*, one the few migratory species in the genus (though four of five subspecies are sedentary and endemic to Pacific Islands), was supported as the first branch in the Australasian clade. Across the nuclear datasets, most of the species-level discordance between analyses centered on the second branch within the clade: whether it was *T. sordidus* (Figs. S8B, 14)*, T. chloris* (Fig. S8A), a sister relationship between the two (Figs. S10, 16), or so poorly supported we considered it a polytomy with the *colonus* clade (Figs. S9, 11–13, 15).

### Mitochondrial discordance and new cases of mtDNA capture within

Todiramphus Compared to nuclear DNA, the mitochondrial data revealed a different picture of the evolutionary history of *Todiramphus* kingfishers in three major ways (Figs. 4, S19). First, the mtDNA-inferred topology matched the ML UCE data in which *pyrrhopygius* diverges prior to the *“leucopygius”* clade (Figs. S9–10, 18). Second, it was the only dataset to show the eastern Polynesian clade (i.e., *T. veneratus* et al.) sister to the rest of the Oceanic clade and Australasian clade. Third, it was the only phylogeny to recover *T. chloris* as the first branch of the Australasian clade and instead recover a significantly different topology from all nuclear datasets: *T. colonus* sister to *T. sanctus* and *T. sordidus*. This relationship has been highlighted before (DeRaad et al. 2023) and was therefore expected. However, we recovered a strongly supported topology of five *T. sacer* taxa embedded within the *T. sanctus* clade, most closely associated with the migratory taxon *T. s. sanctus* and sedentary *T. sanctus macmillani* (endemic to Loyalty Islands, NE of New Caledonia). This group of *T. sacer* taxa with *T. sanctus* mitochondrial DNA included *T. sacer regina* and four allopatric microendemics of Temotu Province of Solomon Islands (*T. sacer ornatus, vicina, brachyurus,* and *utupuae*). Apart from our inability to determine the relationship of *T. s. regina* with nuclear data, these four other Temotu *T. sacer* taxa were either supported as the first few sequentially branching lineages of the *T. sacer* clade (Figs. S9–10, 14–16) or had equivocal support values deeper in the *sacer* complex (Figs. S11, 14–15) across nuclear datasets. In the unrooted phylogenetic network, three of these five taxa were inferred to be early branching and independent lineages on par with splits shown in other species-level taxa (i.e., *T. recurvirostris* and *T. reichenbachii*; Figs. 1C, S17). We had sequenced multiple individuals of all these taxa (n = 2–3 samples) to confirm their curious placement as sequentially branching at the base of the *T. sacer* complex. Yet, we did not find that all individuals of each taxon had *T. sanctus* mtDNA: *T. sacer ornatus* and *T. sacer vicina* had at least one individual in *T. sanctus* and *T. sacer* clades, whereas all samples for *T. sacer brachyurus, regina,* and *utupuae* were recovered within *T. sanctus*.

**Figure 4.**
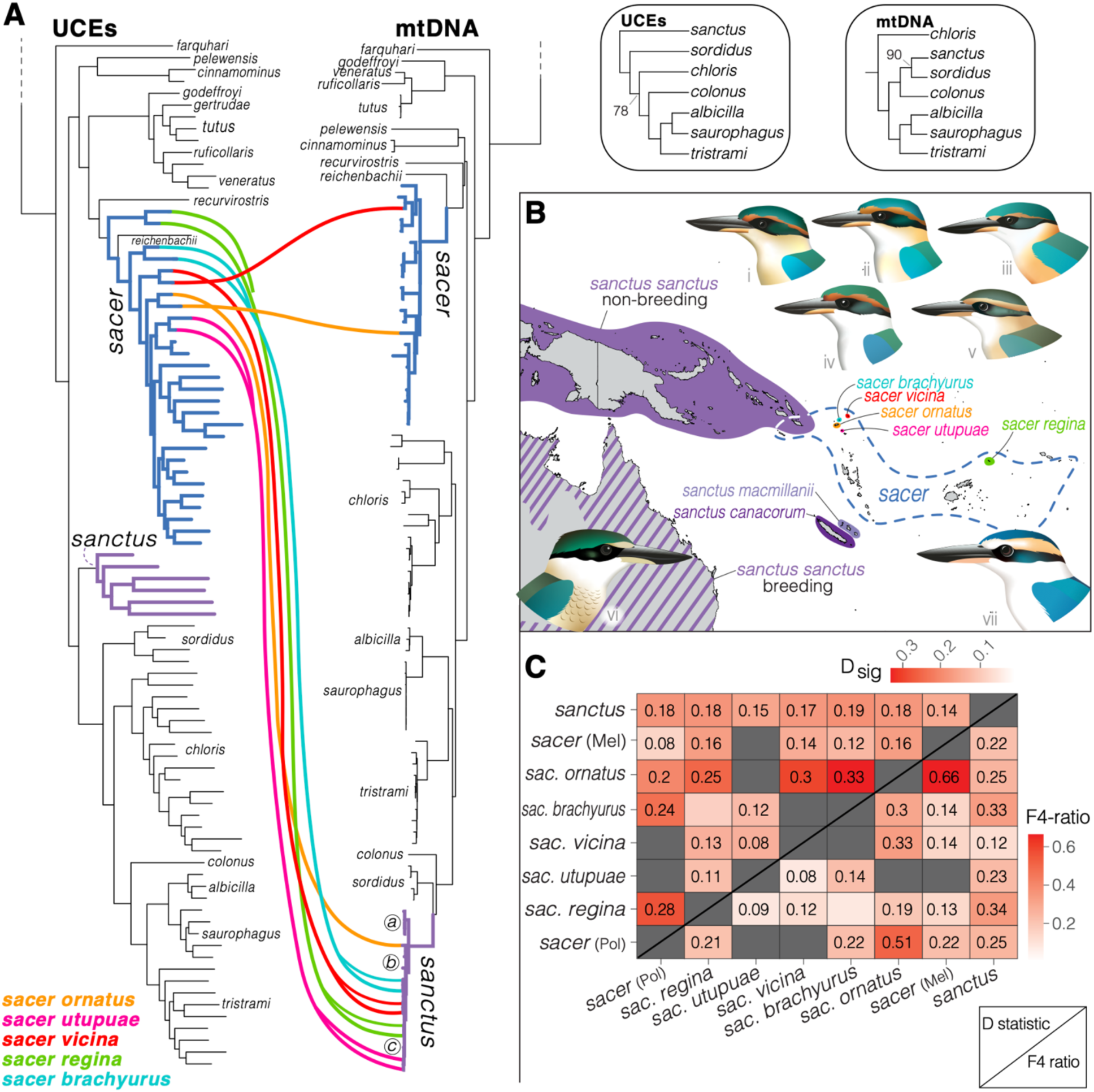
Evidence of gene flow and mitochondrial capture within some Pacific Kingfisher (*T. sacer*) taxa and Sacred Kingfisher (*T. sanctus*). A) Partitioned IQtree phylogenies inferred from the 90% complete UCE matrix and 13 mitochondrial genes. Note that the mtDNA phylogeny for the Australasian clade (*T. sanctus*, *T. tristrami*, and allies) is different from the UCE topology, which has been described previously (DeRaad et al. 2023). In the mtDNA phylogeny, there are two clades of *T. sanctus*: a clade of four of the five sedentary, island endemic *T. sanctus* taxa (a), which is sister to the clade comprising the migratory and nominate taxon *T. sanctus sanctus* (b), the five *T. sacer* taxa (colored lines), and the fifth island endemic sedentary taxon *T. sanctus macmillani* (c). See Fig. S19 for tip labels. B) Range map of both species, highlighting the subspecific ranges of the *T. sacer* taxa with *T. sanctus* mtDNA (see Fig. S1 for the subspecific ranges of all 22 *T. sacer* taxa). Jenna McCullough created the illustrations of kingfisher taxa; names are indicated by roman numerals next to illustrations and are as follows: i, *T. sacer brachyurus*; ii, *T. sacer vicina*; iii, *T. sacer ornatus*; iv, *T. sacer utupuae*; v, *T. sacer regina*; vi, *T. sanctus sanctus*; and vii, *T. sacer sacer* (endemic to Tonga). C) Pairwise results of ABBA-BABA tests from Dsuite, with values in comparisons with FDR-adjusted significant pairwise p values. Above the diagonal line is the most significant (D_min_) for any pairwise comparison and below are F4-ratio values. Only a subset of all comparisons is shown here, see Figs. S20–23 for the complete set.

### Patterns of gene flow

ABBA-BABA tests uncovered biologically plausible scenarios of gene flow across both subsets of data (species-level and the pared down dataset to investigate gene flow between *T. sanctus* and *T. sacer* taxa that have *T. sanctus* mtDNA). Of the 3,544 calculated D-statistics, 858 and 887, respectively, were significant with a FDR-adjusted p value <0.000013. The results show complex patterns highlighting gene flow between *T. sanctus* and mitochondrial discordant lineages, with elevated F4 ratios between *T. sanctus* and *T. colonus* (F4 = 0.34), *T. sordidus* (F4 = 0.31), *T. s. regina* (F4 = 0.34), and the four Temotu *T. sacer* taxa (range of F4 = 0.12–0.33; Figs. S19–20). Though the highest F4 ratios in the pared down dataset are between *T. sacer ornatus* and the Polynesian (F4 = 0.51) and Melanesian (F4 = 0.66) clades of *T. sacer*. With conservative interpretations of these tests by considering the lowest D values between multiple comparisons (D_min_), we still see moderate signals of gene flow between *T. sacer regina* and *T. sanctus* (D_min_ = 0.13; Fig. S21) and *T. sacer* (0.16) in the species-level dataset (Fig. S21). When *T. sacer* taxa that have *T. sanctus* mtDNA are treated as their own tips, we see statistically significant signatures of gene flow between *T. sanctus* and *T. sacer brachyurus* (D_min_ = 0.08; Fig. S19) and between *T. sacer brachyurus* and *T. sacer ornatus* (D_min_ = 0.27). Long swaths of statistically significant and high D-statistic values for every species comparison, such as those shown for *T. colonus,* members of the *leucopygius* and east Polynesian clades (Fig. S21), could reflect either ancient introgression (Fig. S18), incorrect topology, or substitution rate variation caused by bottlenecked populations (Table S4, (Frankel and Ané 2023; Koppetsch et al. 2024).

Network analyses with SNaQ also confirm the presence of hybrid lineages across *Todiramphus* and explain topological discordance across datasets, with the best overall network recovering four hybrid lineages (Fig. S18C). Reticulation events for our best SNaQ topology indicated that between 2–37% of the sampled genome (gamma, γ) was exchanged in each event, with 1-γ being the contribution of the other parental lineage. We inferred an ancient hybridization event that exchanged a sizable portion of the sampled genome between *T. pyrrhopygius* and the ancestor of *T. australasia* + the rest of *Todiramphus* (γ = 0.378). Additionally, we inferred a reticulation event between *T. farquhari* and the ancestor of the east Polynesian clade (γ = 0.236), between *T. chloris* and *T. sordidus* (γ = 0.233), and very low percentage of the sampled genome between *T. sanctus* and the ancestor to the other members of the Australasian clade (γ = 0.022). The other best trees in the two other subsets showed similar patterns of reticulation events with varying levels of γ (Fig. S18). The best inferred networks for other quartet subsets inferred similar and complicated reticulation histories across *Todiramphus*, including branches involving the *T. pyrrhopygius* and the *T. leucopygius* clade, as well as between *T. sanctus*, *T. chloris*, and *T. sordidus*.

### Relationships within species complexes

We found several cases in which *Todiramphus* species complexes do not accurately reflect taxonomic groupings. First, we found that the Talaud Kingfisher (*T. enigma*, endemic to the Talaud islands between Sulawesi and the Philippines), is consistently recovered within the Collared Kingfisher (*T. chloris*) clade. It was most frequently supported as sister to nominate *T. chloris* (*T. c. chloris*, endemic to Wallacea; Figs. 1, 4, S1, S8, S13, S16). Additionally, we consistently found that the two most northern taxa of the *T. sacer* complex, *T. sacer pavuvu* (Pavuvu I.) and *T. sacer mala* (Malita I.) of the central Solomon Islands, were embedded within *T. tristrami* and closely associated with their geographically closest neighbor, *T. tristrami alberti* (main chain of the Solomons Archipelago). We found that the two unnamed *T. sacer* taxa, those from Tucopia Island in the Temotu Province of the Solomon Islands and Anietyum Island in southern Vanuatu, did not group with their assumed subspecific assignment. We never recovered a sister relationship between samples from Tucopia and representatives from its long-assumed subspecies, *T. sacer torresianus,* or between samples from Anietyum and its presumed subspecies, *T. sacer erromangae* (Mayr 1931; Gill et al. 2024; Peters 1945).

## DISCUSSION

As we can efficiently sequence larger and more thoroughly sampled phylogenomic datasets, it is increasingly clear that the evolutionary histories of rapid radiations are complex (Jarvis et al. 2014; Prum et al. 2015; Malinsky et al. 2018; Zink and Vázquez-Miranda 2019; Rota et al. 2022; Mirarab et al. 2024; Stiller et al. 2024). Here, we show that whole genome resequencing of a thoroughly sampled taxonomic dataset of all described forms of a genus of ‘Great Speciators’ has revealed evidence of pervasive gene flow that occurred throughout its rapid radiation. We inferred deep reticulation events as well as more shallow cases of gene flow between both sympatric (*T. sanctus* and *T. sordidus*) and allopatric species (*T. sanctus* and *T. sacer*; Figs. 4, S18–22). This inferred gene flow, in concert with many rapid speciation events that precluded gene coalescence and explains the topological discordance across the four data types used in this study (UCEs, BUSCOs, SNPs, and mtDNA).

Our results confirm past work showing mitochondrial discordance in the Australasian *Todiramphus* clade (DeRaad et al. 2023) and also reveal novel signatures of gene flow with a larger and more comprehensive dataset (Figs. S19–22). Moreover, we uncovered new cases of mitochondrial discordance (Fig. 4) between *T. sanctus* and several *T. sacer* populations that nuclear data inferred as some of the earliest branches within *T. sacer*. Though our limited sampling of a single individual for most taxa precluded us from more detailed population-level analyses, we found signatures of gene flow between *T. sanctus* and Temotu *T. sacer* that have *T. sanctus* mtDNA (Figs. S19–22). The inference of these hybrid lineages as divergent and non-monophyletic lineages suggests that they follow the ‘artefactual branch effect’ (Chan et al. 2020b, 2023; Pyron et al. 2022). Our results suggest that there are numerous violations of the traditional bifurcating species tree model within *Todiramphus* kingfishers. A next step towards understanding interspecies relationships would be a sliding window approach to determine if particular genomic regions are disproportionately driving the levels of discordance observed here, as has been shown in early branches of Neoaves (Mirarab et al. 2024; Stiller et al. 2024).

### Gene flow within Todiramphus kingfishers

Studies that employ genome-wide data are increasingly demonstrating the prevalence of gene flow during periods of rapid diversification (Edelman et al. 2019; Taylor and Larson 2019; Guo et al. 2022; Ocampo et al. 2022; Jensen et al. 2023; Jiang et al. 2024; Stiller et al. 2024; Zhao et al. 2024). Here, we uncovered novel patterns of gene flow between *Todiramphus* kingfishers as they rapidly diversified over the last 10 Ma. Recent population-level studies that employed either mitochondrial (O’Connell et al. 2019) or RAD-seq data (DeRaad et al. 2023) did not find signatures of gene flow between *T. sanctus* and other members of the Australasian clade (*T. chloris*, *T. sordidus*, *T. colonus*, etc.). Here, comparisons between these lineages showed relatively lower signatures of gene flow than other pairwise comparisons, such as between *T. sanctus* and *T. sacer,* and within the *T. sacer* complex (Figs. S20–23). Network analyses inferred a very low percentage of the sampled genome (2.2%) shared between *T. sanctus* and the ancestor to the remaining members of the Australasian clade (Fig. S18). This novel inference of gene flow was likely missed in prior studies due to the ever-growing size and scope of molecular datasets brought to bear on this group (three mtDNA genes for O’Connell et al. 2019 vs. ∼2,000 SNPs for Deraad et al. 2023 vs. >85K SNPs and thousands of gene trees for this study). We are not the first to uncover gene flow within Indo-Pacific kingfishers. Studies that employ phylogenomic data showed persistent gene flow but maintenance of species-boundaries between the two sympatric *Syma* species (sister genus to *Todiramphus*) across an elevational gradient in New Guinea (Linck et al. 2020) as well as across *Ceyx,* the other ‘Great Speciator’ kingfisher genus (Lim et al. 2010; Andersen et al. 2013; Shakya et al. 2023). Overall, our study fits into a larger picture of gene flow between island endemic avian lineages, including within Fijian groups such as honeyeaters and bush-warblers (Gyllenhaal et al. 2020; Mapel et al. 2021) and Indonesian monarch flycatchers (Andersen et al. 2021).

Many of the contemporary patterns of gene flow that we observed in this study involve one species in particular, the Sacred Kingfisher *T. sanctus*. Because we had limited sampling per taxon, we could not pursue more detailed tests of gene flow between *T. sanctus* and other species, such as Admixture (Alexander et al. 2009) or fastsimcoal2 (Excoffier et al. 2021). However, observed mitochondrial discordance of five *T. sacer* taxa (*regina*, *ornatus, utupuae*, *vicina*, and *brachyurus;* Fig. 4) and elevated estimates of D_min_ between *T. sanctus* and *T. sacer* from ABBA-BABA tests (Figs. S19, S21) confirms the presence of gene flow. Moreover, one of these *T. sacer* lineages (*regina*) is fully supported in the *T. sanctus* clade in one phylogeny (BUSCO IQtree; Fig. S16). *Todiramphus sanctus* is notable because it is one of two truly migratory *Todiramphus* species alongside the Forest Kingfisher (*T. macleayii*; Woodall 2001). Moreover, only the nominate subspecies of *T. sanctus* is known to be migratory—the other four taxa within this complex are considered sedentary and endemic to separate Pacific islands (Fig. S1). We inferred gene flow between *T. sanctus* and Torresian Kingfisher (*T. sordidus*), a species that also breeds in Australia. However, during its non-breeding season, *T. sanctus* occupies a wide swath of the Indo-Pacific: from as far west as Sumatra, across Wallacea, New Guinea, and northern Melanesia, and as far east as Makira in the Solomon Islands (Fry et al. 1992; Woodall 2001). Therefore, in the non-breeding season, *T. sanctus* is sympatric with upwards of 42 *Todiramphus* taxa (representing 15 species), compared to only five taxa (3 species) in the breeding season. However, the *T. sacer* taxa that we found to have *T. sanctus* mtDNA are not sympatric with wintering *T. sanctus*, as there are no records of *T. sanctus* in Temotu Province, Solomon Islands, or even as far east as Wallis and Futuna (*T. sacer regina*; see Appendix 2 for more details on breeding records of *T. sanctus*). This suggests a complicated history of island colonization, historical gene flow, and range contraction between these two species.

With the lack of substantial breeding records of *T. sanctus* outside of Australia, we are left with one central question: why do we infer gene flow between *T. sanctus* and Temotu *T. sacer* (*ornatus, utupuae*, *vicina*, and *brachyurus)* and *T. s. regina* (Fig. 4) when we do not have sympatric records (breeding or otherwise) of these taxa? Notably, the only islands that sedentary *T. sanctus* inhabit are conspicuously void of *T. sacer*. Geographically, the closest breeding *T. sanctus* taxon to an island with *T. sacer* is *T. sanctus macmillani* (Loyalty Islands) and the undescribed *T. sacer* population on Anietyum [also written as Anatom], approximately ∼220 km apart. *Todiramphus sanctus macmillani* is notable because it was placed in the mitochondrial clade with the migratory *T. sanctus sanctus* and all five *T. sacer* (Fig. 4). This pattern of non-overlapping breeding ranges of sedentary *T. sanctus* and *T. sacer* parallels a pattern described by Mayr (1932b) for Fijian whistlers (*Pachycephala*) in which the archipelago was first colonized by one group and then replaced by another, forming hybrid populations as the second wave of colonists swept through the archipelago. Notably, crown *T. sanctus* predates that for *T. sacer* (McCullough et al. 2019, McCullough et al. *in prep*), so perhaps the ancestral range of *T. sanctus* had more widespread sedentary populations across the South Pacific prior to members of the Oceanic clade’s back colonization from Eastern Polynesia towards Melanesia. This hypothesis needs further testing, but it would explain the evidence of hybridization (Fig. 4) without the contemporary presence of *T. sanctus* in the areas where *T. sacer* lineages possess *T. sanctus* mtDNA.

### Other sources of topological discordance

In addition to the demonstrated gene flow across the *Todiramphus* radiation (Fig. 4), topological discordance could also be attributed to other biological and methodological factors. Incomplete lineage sorting (ILS) is a well-known and pervasive source of topological discordance, especially within rapid radiations (Suh et al. 2015; Lopes et al. 2020). However, teasing apart the relative contribution of gene flow versus ILS is difficult. DeRaad et al. (2023) suggested that mitonuclear discordance within the Australasian *Todiramphus* clade was driven largely by ILS rather than nuclear gene flow or mitochondrial capture, based on largely non-significant pairwise comparisons of the ABBA-BABBA D-statistic. Here, gene and site concordance factors (gCF/sCF) between major *Todiramphus* topologies similarly suggest that ILS is likely a significant source of topological conflict across the entire genus (Fig. 3). However, ancient reticulation events consistently inferred in similar areas of the tree (Fig S18) and significant pairwise D-statistic comparisons using our comprehensive SNP dataset (Figs. S20–23), together suggest that ongoing gene flow during diversification contributed to signals of genome-wide discordance in the Austalasian clade (Figs. 2–3). While DeRaad et al. (2023) rejected mitochondrial capture within the Australasian *Todiramphus* clade, our expanded taxonomic sampling across the genus reveals clear cases of mitochondrial capture of recently derived *T. sanctus* haplotypes in multiple *T. sacer* lineages, highlighting the overall complexity of the heterogeneous evolutionary processes simultaneously contributing to generate the patterns of relatedness we observe among extant *Todiramphus* lineages.

Alternatively, methodological explanations like the source (tissue or toepad) or type of DNA (exonic vs. intronic) could also explain some of the topological discordance among data types and tree building methods. Because our dataset relied heavily on toepad samples, we took additional steps at all levels of this study to ameliorate the effects of degraded DNA on topological inference. In addition to special wet lab processing of toepad-sourced genomes, we performed additional UCE alignment cleaning (Smith et al. 2020) and novel methods of *in silico* BUSCO extractions (see Appendix 2). Moreover, we explored filtering approaches for MSC tree building methods. ASTRAL is known to perform poorly when input gene trees are built from short alignments of uninformative loci (Mirarab 2023), a pervasive issue for datasets utilizing degraded DNA samples from museum specimens (Andersen et al. 2019; Salter et al. 2019, 2022; Smith et al. 2020; Roycroft et al. 2021; Huynh et al. 2023). For datasets that have a high degree of ancient or historical samples, there is often an artefactual pattern in which high quality and historical samples form distinct clusters (Fig. S2) rather than being uniformly distributed in the species tree. Before filtering for parsimony informative (PI) sites, ASTRAL-inferred species trees showed a clear artefactual pattern of toepad-sourced samples clustering at the base of their respective species-complexes (Figs. S2, S4). This pattern disappeared when we analyzed smaller sets of gene trees based on alignments with more PI sites (Figs. S11, S15). Although filtering based on informativeness can introduce bias (Musher et al. 2019; Burbrink et al. 2020), the approach has been used with success elsewhere (e.g., Chan et al. 2020a; Leite et al. 2021; Forthman et al. 2022). We preferred it here because it is a general approach to deal with these issues rather than clade-specific filtering approaches (Chen et al. 2018) or by dropping important tips represented by toepads. Additionally, the clades that represented the most topological discordance were from those represented by high-quality tissues (i.e., *T. farquhari*, *T. pyrrhopygius*) or were in clades with relatively few toepad-sourced samples (i.e., the *leucopygius* clade, *T. sordidus*; Fig. 2).

The type of DNA from which molecular markers are derived (i.e., exonic vs. intronic regions) could also contribute to conflicting topological patterns. UCEs are largely sourced from intronic and non-coding regions of the genome whereas BUSCOs are all exonic markers from coding regions. The difference in selective pressures and therefore rates of fixed mutations, has been shown to have a substantial impact in the overall topology inferred by these markers, such as in higher-level phylogenomic studies of all birds (Jarvis et al. 2014; Prum et al. 2015; Reddy et al. 2017). Yet, we generally find that these data types produce largely similar topologies (Figs. 2–3). Though less commonly used in species-level relationships of birds, these molecular markers are more widely used to infer phylogenies outside of birds, such as in plants (Timilsena et al. 2022; Walden and Schranz 2023), fungi (Kanzi et al. 2020), arthropods (Dias et al. 2020; Van Damme et al. 2022).

### Taxonomic recommendations

In line with a growing body of work that has uncovered unsuspected species-level diversity in geographic radiations across the Indo-Pacific (Andersen et al. 2013; Reeve et al. 2023; Pedersen et al. 2018; Irestedt et al. 2013), we too found that some taxa in our dataset require taxonomic revisions based on their novel systematic relationships (see Appendix 2 for further discussion). Roughly half of the *Todiramphus* diversity we sampled has never been included in a molecular phylogenetic analysis. Many newly added taxa fall within their presumed species groups, such as all of the subspecific diversity within *T. tristrami* (Fig. 1), but there are some instances that require taxonomic changes. When Ernst Mayr described geographic variation of kingfishers across the Pacific, he noted that two populations on Tucopia (Temotu Province, Solomon Islands) and Aneityum (Tafea Province, Vanuatu) islands were each phenotypically distinct from those on nearby islands, but he refrained from naming these as subspecies (Mayr 1931b). He wrote of the birds from Anietyum, they were “indistinguishable from typical *juliae…* only the distribution keeps me, for the present, from uniting it immediately with *juliae*” (Mayr 1931b, pg 4). To our eyes, Aneityum *sacer* are more similar to those in central Vanuatu (*T. sacer juliae*) with white underparts and collar rather than the buffy *T. s. tannensis* on Tanna Island. However, instead of considering this a separate taxon or grouping this population with the closest island (Tanna), Aneityum birds have long been considered as part of *T. sacer erromangae*, forming a curious leapfrog distribution of white-bellied birds (*erromangae* and the Aneityum form) sandwiching buffy-bellied birds (Tanna). We would expect that if Aneityum birds are indeed part of *T. s. erromangae* or *tannensis*, our genetic samples would be closely associated with one of these taxa. For the samples from Tucopia, Mayr refrained from naming these birds as a subspecific taxon because their plumages were “too worn” (Mayr 1931b), and they have been considered either a part of *T. s. torresianus* (Banks Is., Vanuatu) or *T. s. melanodera* (Vanikolo in Temotu Province). Whereas we sometimes found Aneityum *sacer* grouping in a subclade with *T. s. erromangae* or *tannensis* and Tucopia *sacer* (Fig. 1), we never recovered Aneityum *sacer* or Tucopia *sacer* as sister to either of their presumed subspecies. We suspect these could represent independent lineages from their long-presumed subspecies but we defer naming these two populations as subspecies at this time.

## CONCLUSIONS

Geographic radiations across the Indo-Pacific have been critical to the development of influential speciation theories, with genome-scale datasets providing new and accelerated insights into the complicated evolutionary histories of such rapidly diversifying clades. In this study we show that whole genome resequencing and complete sampling of a rapid radiation of mostly island kingfishers does not outright resolve the evolutionary relationships of the group. Instead, we found that four types of data (UCEs, BUSCOs, SNPs, and mtDNA) and tree building methods (ML, MSC, networks) did not infer a single well-supported and concordant species-level topology. Phylogenetic network analyses suggest that ancient introgression is likely the cause of these topological differences. The discovery of several *T. sacer* populations falling in the mitochondrial clade of a widespread and migratory taxon, *T. sanctus*, and the inference of gene flow between them ABBA-BABA tests, suggests a potential historical hybrid zone between these two species as islands were successively colonized. We were only able to pick up on these signals of hybridization because we thoroughly sampled all described diversity within the genus by relying on a sizable number of toepad-sourced data. Future studies that investigate rapid radiations in this region should incorporate widespread taxonomic and population level sampling, while also sampling large amounts of genetic data to tease apart the interacting effects of gene flow and incomplete lineage sorting on complex evolutionary histories.

## ACKNOWLEDGEMENTS

This study unequivocally would not have been possible if not for the countless people since 1887 who have dedicated their time to collecting, preparing, preserving, and loaning bird specimens and their associated tissues to the world’s scientists. Nearly 30% of our sampling was from the American Museum of Natural History’s Whitney South Seas Expedition (1920–1941), therefore we would like to acknowledge some of its most prolific collectors: Hannibal Hamlin, Rollo H. Beck, William F. Coultas, and Ernst Mayr. We specifically thank the curators and collections managers of the American Museum of Natural History, Australian National Wildlife Collection, Charles R. Conner Museum, Delaware Museum of Natural History, the Field Museum, University of Kansas Biodiversity Institute and Natural History Museum, the Natural History Museum in London, Natural History Museum of Los Angeles County, Louisiana State University Museum of Natural Science, Museum of Comparative Zoology, Natural History Museum of Geneva, Museum of Southwestern Biology, Museum of Vertebrate Zoology, New York State Museum, Smithsonian National Zoological Park, University of Michigan Museum of Zoology, United States National Museum, University of Washington Burke Museum, Western Australia Museum, and Yale Peabody Museum. We thank Devon DeRaad, Ethan Gyllenhaal, David Tan, Fernando Machado-Stredel, Rob Lanfear, Laura Kubatko, David Bryant, Leo Joseph, and Nathan Kolbow for helpful discussions on the analyses presented in this manuscript. We would like to thank the UNM Center for Advanced Research Computing, supported in part by the National Science Foundation (NSF), for providing the research computing resources used in this work. This study was funded by NSF (DEB 2112467 to C.E.M., S.J.H., and M.J.A) and the following sources to J.M.M.: the American Museum of Natural History Frank Chapman Research Grant, UNM Biology Department Alvin R. and Caroline G. Grove Research and Richard B. Forbes Conservation awards, British Ornithologists’ Union Small Ornithological Research Grant, and the American Ornithological Society Werner and Hildegard Hesse Research award.

## DATA AVAILABILITY

Supplementary material, including alignments, tree figures, and other information (supplementary methods, discussion, figures, and tables) are available in the Dryad data repository (available upon acceptance). Raw read data sequenced for this project is available on Genbank (accession upon acceptance).

## APPENDICES

Appendix 1: Combined excel spreadsheet that includes five supplementary tables.

Appendix 2: Combined PDF of supplementary methods, discussion, and figures.

