## Appendix 2 for "Phylogenomics of a genus of ‘Great Speciators’ reveals rampant incomplete lineage sorting, gene flow, and mitochondrial capture in island systems"

**Appendix 2:** Supplementary Information for **"**Phylogenomics of a genus of 'Great Speciators' reveals rampant incomplete lineage sorting, gene flow, and mitochondrial capture in island systems"

Jenna M. McCullough, Chad Eliason, Shannon Hackett, Corinne E. Myers, and Michael J. Andersen

<u>The following Supplementary Information includes:</u>

Supplementary Methods and Discussion

Supplementary figures 1–23

**Supplementary Methods**

*Sampling and laboratory methods*

For tissue-sourced samples, we used the Qiagen DNeasy kit to extract genomic DNA. For toepad-sourced samples, we performed phenol-chloroform extractions with the addition of Dithiothreitol (DTT) and gel phase-lock tubes (Murphy and Hellwig 1996) to increase yields for toepad samples, following best practices (Billerman and Walsh 2019; Tsai et al. 2019). We estimated fragment length with gel electrophoresis for tissue-sourced samples and sheared 250 ng of genomic DNA, aiming for 500 base-pair (bp) fragments, with a Covaris M220 focused-ultrasonicator (50 W peak incident power, 10% duty factor, and 200 cycles per burst for 55–75 s, dependent on sample quality). Toepad-sourced extractions are known to have much smaller average fragment sizes (Tsai et al. 2019) and therefore do not need to be sheared further if older than 30 years. Instead, we performed a Uracil-damage treatment (Rohland et al. 2015) with USER enzyme (New England Biolabs) to remove deaminated cytosines from all toepad-sourced extractions prior to library preparation. This method reduces natural DNA damage patterns and has been used for other projects reliant on whole-genome resequencing of toepad-sourced samples (Irestedt et al. 2022). We quantified DNA extractions with a Qubit 3.0 Fluorometer (ThermoFisher Scientific) for all samples prior to library preparation.

We prepared Illumina libraries with half the volume per sample using the Kapa Biosystems Hyper Prep Kit and the iTruStub dual-indexing system (baddna.org). For toepad-sourced samples, we did not sonicate prior to library prep, used Eppendorf Lo-Bind tubes for all non-thermocycler reactions, doubled ligation time (30 min), and increased concentrations of all AMPure (Beckman Coulter) bead cleanups to 3X sample volume. For all samples, we used the Qiagen GeneRead Size Selection Kit to remove fragments <150 bp prior to sequencing and a Bioanalyzer to visualize fragment sizes. We pooled toepad- and tissue-sourced libraries separately (10–15 toepad libraries per lane and 15–16 tissue libraries per lane) and sequenced them across 12 S4 PE-150 lanes of an Illumina NovaSeq 6000 at the Oklahoma Medical Research Foundation.

*Removal of "dirty ends" of UCE loci*

During preliminary phylogenetic exploration, we determined that many of the toepad-sourced samples had exceptionally long branches (see below for phylogenetic analyses and Table S1 for these samples). Biologically unrealistic long branches such as these can contribute to long-branch attraction and therefore incorrect phylogenetic relationships (Felsenstein 1978); this issue has been shown as a byproduct of poor alignment trimming contributing to "dirty ends" of UCE loci sourced from historical samples, including those sourced from avian toepads (McCormack et al. 2015; Smith et al. 2020; Salter et al. 2022). We removed these artifactual "dirty ends" following the bioinformatic pipeline by Smith et al. (2020). To do so, we chose clade-specific reference samples to which we aligned trimmomatic cleaned reads; to identify these reference samples, we expanded fasta files (with "phyluce_assembly_explode_get_fastas _file") and chose a closely related, tissue-sourced sample with the highest number of UCE loci based on initial maximum likelihood analyses (Table S2). We used bwa and SAMtools to index the UCE contigs of reference samples and align cleaned reads; we removed low-quality data in the flanking regions by dropping sites with <5X coverage and quality scores of <20. We manually incorporated these cleaned samples back into the phyluce pipeline by adding the nucleotide data into a combined unaligned fasta file (with all other non-cleaned samples) and again aligned and trimmed loci with the same settings as above. Our final UCE dataset was a 90% complete matrix comprising multiple individuals for some taxa that had at least 105 of 117 samples present at each UCE locus (Table S3).

*Filtering of UCE loci for ASTRAL*

We used the summary-statistic method ASTRAL-III v.1.15.2.4 (Zhang et al. 2018; Rabiee et al. 2019) to estimate the species tree. This summary-statistic MSC method is sensitive to fragmentary data and depends on high quality gene trees (Mirarab 2023). Because toepad-sourced samples often have a higher degree of missing data within individual alignments, we filtered the 90% complete matrix to drop samples from individual alignments if they comprised more than 25% of ambiguous nucleotides (N) or 70% of gaps. We performed model selection and assessed gene tree support with 1,000 ultrafast bootstraps on alignments. We further filtered our gene trees based on PI sites (determined with phyluce_align_get_ informative_sites). With a threshold of >50 PI sites (1583 gene trees), our toepad-sourced samples showed a clear artifactual pattern in which they clustered at the base of clades to the exclusion of tissue-sourced genomes (Fig. S2), a known issue for ASTRAL that is caused by poor quality and/or uninformative gene trees (Mirarab 2023) and the inclusion of DNA from degraded sources

(Smith et al. 2020). Therefore we further filtered gene trees based on loci with at least 100 PI sites, which yielded 125 highly informative gene trees. For ASTRAL, we increased the number of searches (-r) and subsampling (-s) from the default of 4 to 8. To estimate a concatenated ML phylogeny of the 90% complete matrix, we used RaxML-NG v.1.2.0 (Kozlov et al. 2019), assuming a general time-reversible model of nucleotide substitution and gamma-distributed rates among all sites (GTR+G) and evaluated support with the autoMRE function (set to 1000 BS). Additionally, we performed a partitioned analysis with IQtree 2.1.4 (Chernomor et al. 2016; Minh et al. 2020), implementing modelfinder and assessing support with 1,000 replicates of Ultrafast bootstraps (Hoang et al. 2018).

### Supplementary Discussion

*Gene flow within* Todiramphus *kingfishers*

Prior to this study, there were two pieces of evidence that the migratory *T. sanctus* breeds outside of Australia. The first is a very brief record (French 1957) that only lists a clutch size of three eggs for *T. sanctus* on the Three Sisters Islands (small islands off the northern coast of Makira, Solomon Islands). However, this record lists three breeding *Todiramphus* (at the time was considered "*Halcyon*"): *T. saurophagus*, *T. sanctus*, and *T. chloris.* Confusingly, at the time, "*Halcyon chloris*" could have been two different taxa that are considered separate species today: *T. tristrami alberti* (widespread across the Solomon Islands) or *T. sacer sororum* (endemic to just the Three Sisters (Woodall 2001; Gill et al. 2024). *T. sacer sororum* appears similar to typical *T. sacer*: white underparts, white collar, and a sexually dichromatic superciliary stripe (hereafter referred as "eyebrow") that extends over the eye and connects around the back of its head, a feature common among *T. sacer* taxa (females are white and males are rufous). *T. tristrami alberti* has dark buffy underparts and no distinctive eyebrow stripe (matching the green-ish blue of the crown), which is much more similar to *T. sanctus*' light buff underparts and no distinctive eyebrow. In the field, *T. t. alberti* could easily be mistaken for the migratory *T. sanctus*. Therefore, it is more likely a case of mistaken field ID rather than a confirmed breeding record outside of Australia. The second record is from the first phylogeographic study of the genus (Andersen et al. 2015b): two birds from Nendo Island were inferred with mtDNA to be in separate clades, with one in *T. sanctus* (KU tissue #19403) and the other with *T. sacer* (KU

19404). Because of its placement in the *T. sanctus* clade, the individual was considered a misidentified *T. sanctus* rather than *T. sacer ornatus* with *T. sanctus* mtDNA. However, comparisons of the specimen in question confirm it is not a case of misidentification because plumage characters match *T. sacer ornatus* (a male with a rufous eyebrow and unspotted rufous breast matching other *T. sacer ornatus* collected during the Whitney South Seas expedition). We sequenced the "*T. sanctus*" from Nendo in this study (KU 19403; Table S1) and found that all nuclear data and methods supported its position within *T. sacer* but mitochondrial data placed it again in *T. sanctus* (Fig. 4). However, the mitochondrial capture of *T. sanctus* mtDNA is not complete, as evidenced by both Andersen et al. 2015 and our finding that a toepad-sourced sample from the 1920's also inferred with other *T. sacer.*

*Other sources of topological discordance*

Another step taken to decrease the methodological influence of data source dealt with the extraction of BUSCO loci from toepad-sourced genomes. We found that we recovered poor quality BUSCO data when we extracted these markers from toepad-sourced genomes. This pattern was easily distinguishable because they clustered together in phylogenetic analyses in a non-biological pattern, similar to the situation noted above (Fig. S3). When we instead extracted BUSCO loci from coordinates from loci from our reference genome, this artifactual pattern disappeared and toepad-sourced genomes no longer clustered together (Figs. S15–16). Though more often used to determine the general completeness of a genome (Simão et al. 2015; Seppey et al. 2019; Feng et al. 2021), BUSCO loci are less frequently used as molecular markers in avian phylogenetics (but see Alaei Kakhki et al. 2023; Musher et al. 2024). Relatively few studies, to our knowledge, have extracted BUSCO loci from genomes sourced from degraded DNA for use in phylogenetic inferences (Musher et al. 2024). Considering the exponential increase in phylogenomic studies that include degraded DNA sourced from museum samples ("museumomics"; Raxworthy and Smith 2021; Ernst et al. 2022), future studies that aim to incorporate BUSCOs from degraded sources may want to explore alternate methods of extraction like those in this study.

*Taxonomic recommendations*

We found that *T. enigma*, an endemic of the Talaud Islands, Indonesia, is closely related to nominate *T. chloris*, often to the exclusion of the rest of the *T. chloris* taxa (Figs. 1, S8–10, S15–17). Though originally described as a distinct species (Hartert 1904), *T. enigma* was long considered a subspecies of *T. chloris* until it was split based on phenotypic and life history differences (Riley et al. 1998). Since its elevation to species status, there have been relatively few studies on its ecology and no further information on the breeding biology of *T. enigma* or *T. chloris* on Talaud (Riley 2003; Kelly et al. 2017). *Todiramphus enigma* was not sampled by Andersen et al. (Andersen et al. 2015)(2015b) and higher-level analyses that included *T. enigma* lacked sufficient sampling to place this taxon unequivocally (Andersen et al. 2018; McCullough et al. 2019). Our results suggest that *T. enigma* is best treated as part of *T. chloris*; however, a more nuanced explanation may involve undetected gene flow between these sympatric species or due to a toepad artifact, since our sample of *T. enigma* was sourced from a specimen from 1886 with 6.8x average coverage. Considering that *T. enigma* is a near-threatened species endemic to three small islands (~1,250 $km^2$ total land area), it is susceptible to threats from ongoing deforestation. More focused research is needed before lumping *T. enigma* with *T. chloris*.

We consistently found that two *T. sacer* taxa were embedded within the *T. tristrami* clade (Figs. 1, S8–17). *Todiramphus sacer pavuvu* and *mala* are some of the most northern representatives of *T. sacer*, endemic to Pavuvu (also referred to as the Russell Islands) and Malaita, respectively, in the Solomon Islands. *Todiramphus tristrami* and *T. sacer* were, among other *Todiramphus* species (*T. sordidus, T. saurophagus*, *T. colonus*, *T. albicilla*), part of the *T. chloris* species complex when it was thought to comprise 49 subspecies prior to Andersen et al. (2015). In that study, neither of these taxa were sampled and their species identification was likely based on geographic range and plumage characters. Here, *T. sacer pavuvu* and *mala* were sourced from toepads that were inferred to have a close association with *T. tristrami alberti*, which is found across the main Solomons Archipelago. Morphologically, these two taxa are quite similar to *T. tristrami alberti*, with their original species descriptions only highlighting minute differences from *alberti* (Mayr 1935). Considering their non-monophyly with other *T. sacer*, we recommend treating *pavuvu* and *mala* within *T. tristrami*. Whether or not *pavuvu* and *mala* are best treated as valid subspecies remains an outstanding question. To our knowledge, *mala* is the only *T. tristrami* taxon on Malaita, an island that has several other endemic bird taxa, so it seems reasonable to treat *mala* as different from *alberti*. The question of *pavuvu* is a little more uncertain because we do not know if true *alberti* occurs on the Pavuvu Islands. It is unlikely that *alberti* and *pavuvu* co-occur on these tiny islands, but until this point can be resolved, we recommend maintaining *pavuvu* as a subspecies within *T. tristrami*.

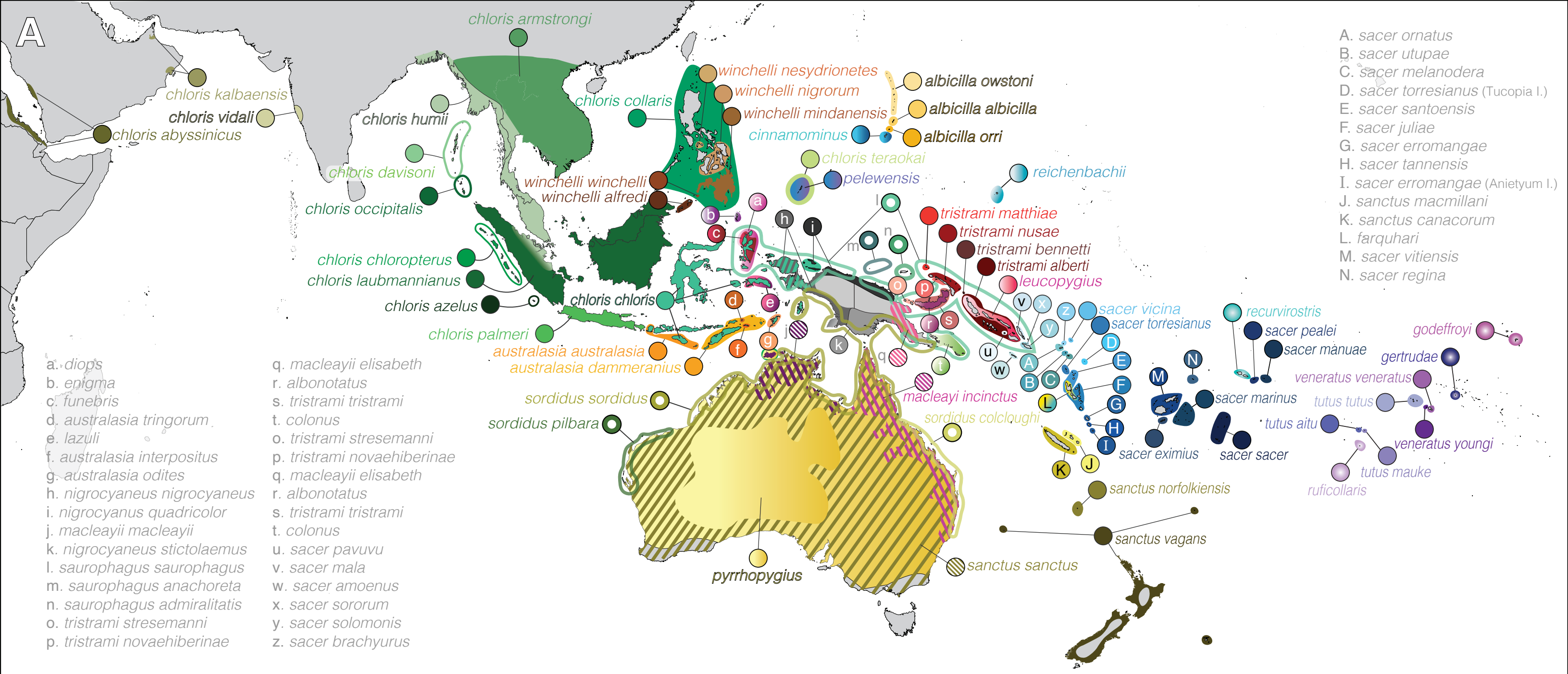


Figure S1. Map of *Todiramphus* kingfishers. Taxon-specific ranges of 93 taxa within the genus, with diagonal lines showing sympatry in some cases.

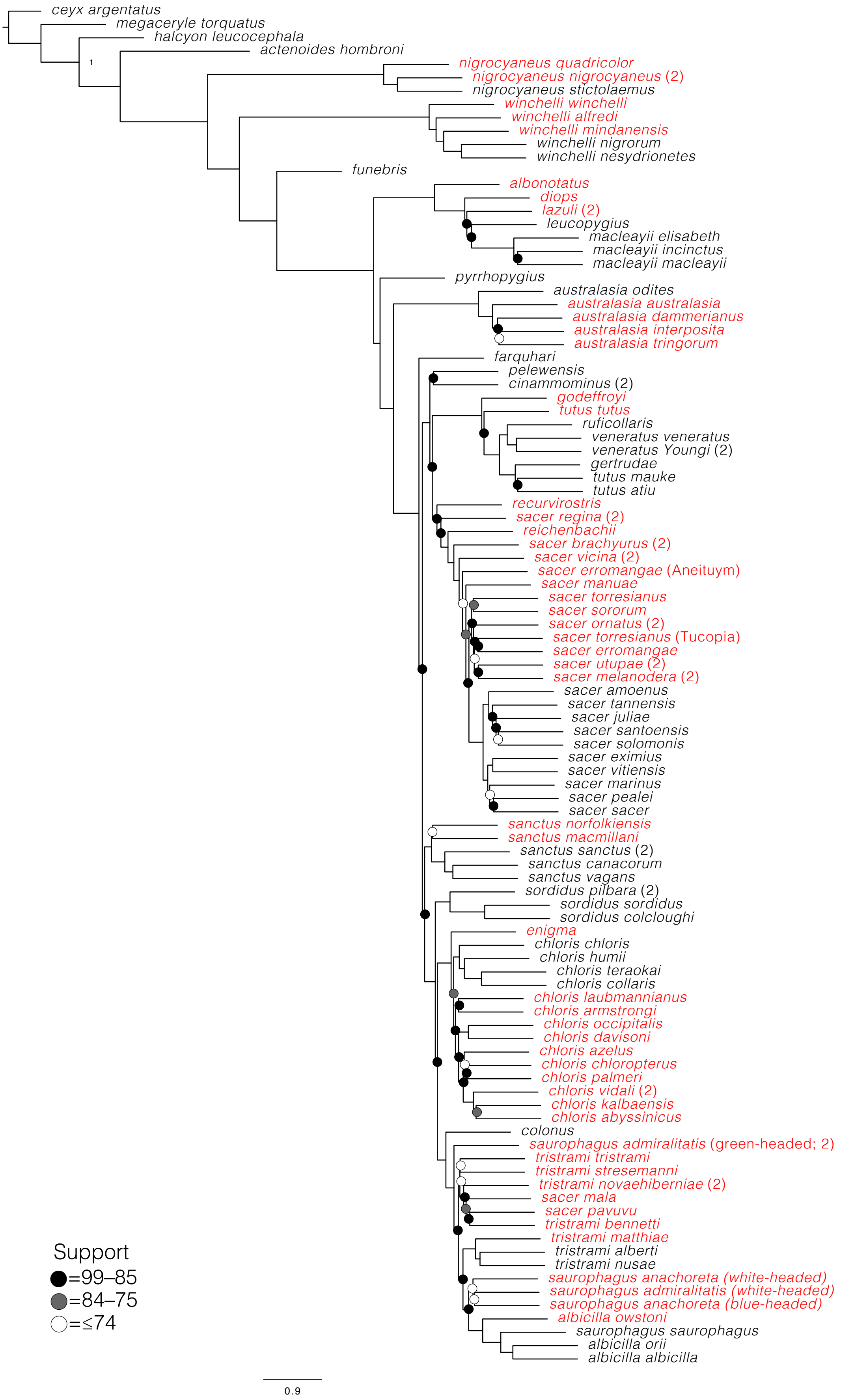


Figure S2: ASTRAL phylogeny of UCE 90% complete matrix with a threshold of >50 Parsimony informative sites (1583 gene trees). Red names are toepad-sourced samples and the following number in parenthesis indicates how many samples are represented in that tip that were separate in gene trees (lumped by taxon).

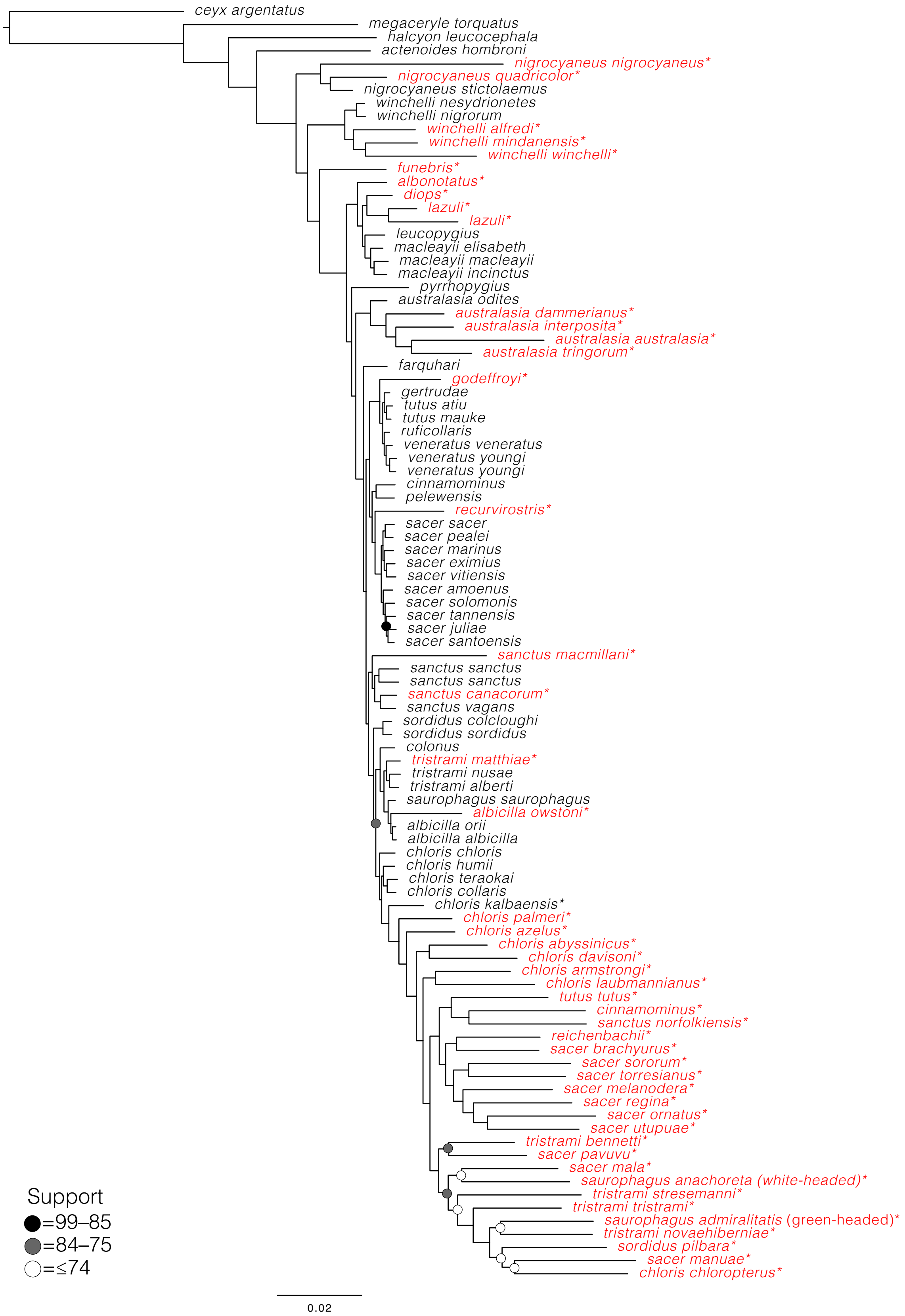


Figure S3: IQtree phylogeny of 8,012 BUSCO loci when extracted from genomes using the BUSCO program. Toepad-sourced samples are red with an asterisk (*). Note the artificially long branches and artefactual clustering toepad samples.

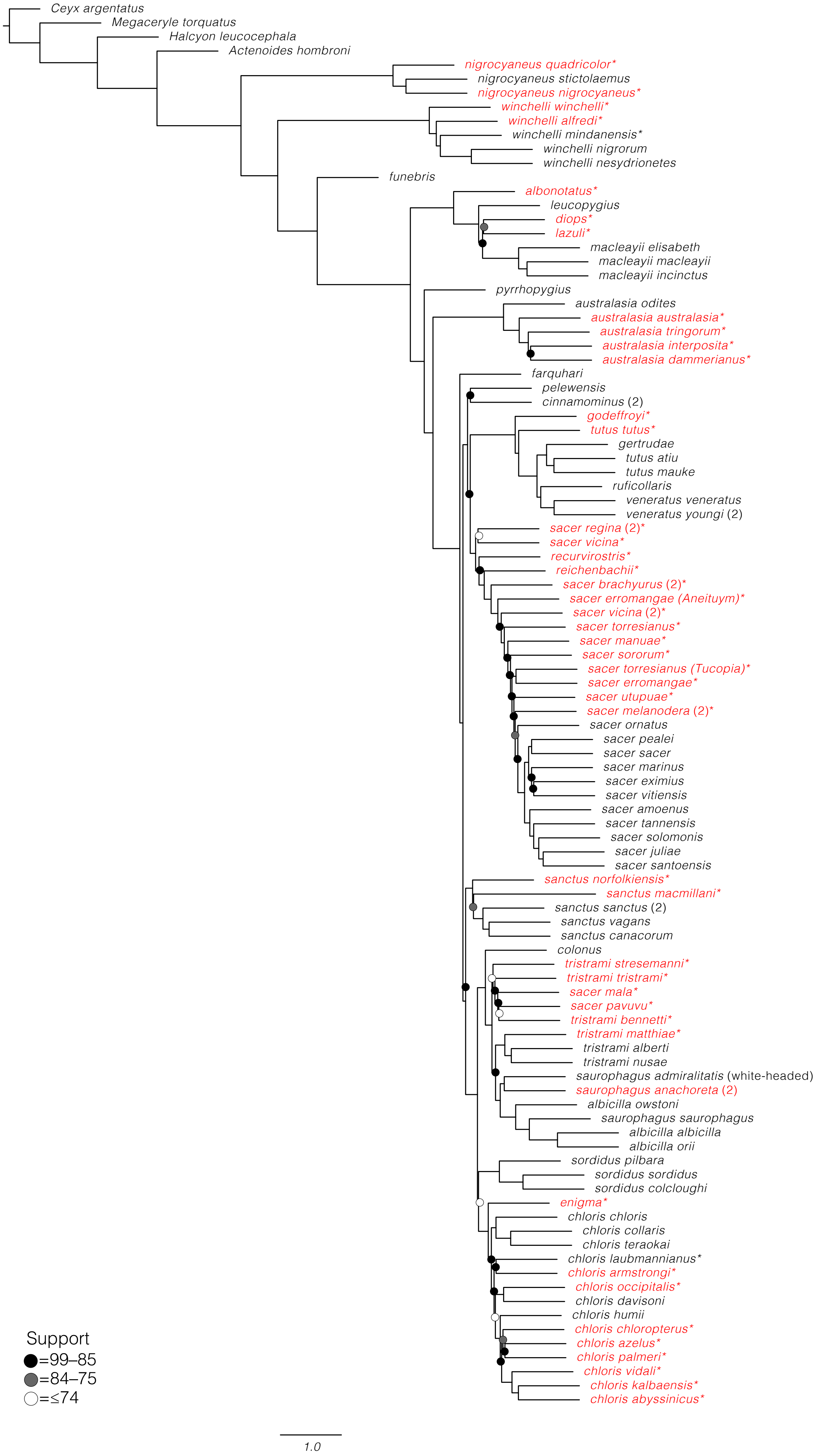


Figure S4: ASTRAL phylogeny of BUSCO loci with a threshold of >20 Parsimony informative sites (4,177 gene trees). Red names are toepad-sourced samples and the following number in parenthesis indicates how many samples are represented in that tip that were separate in gene trees (lumped by taxon).

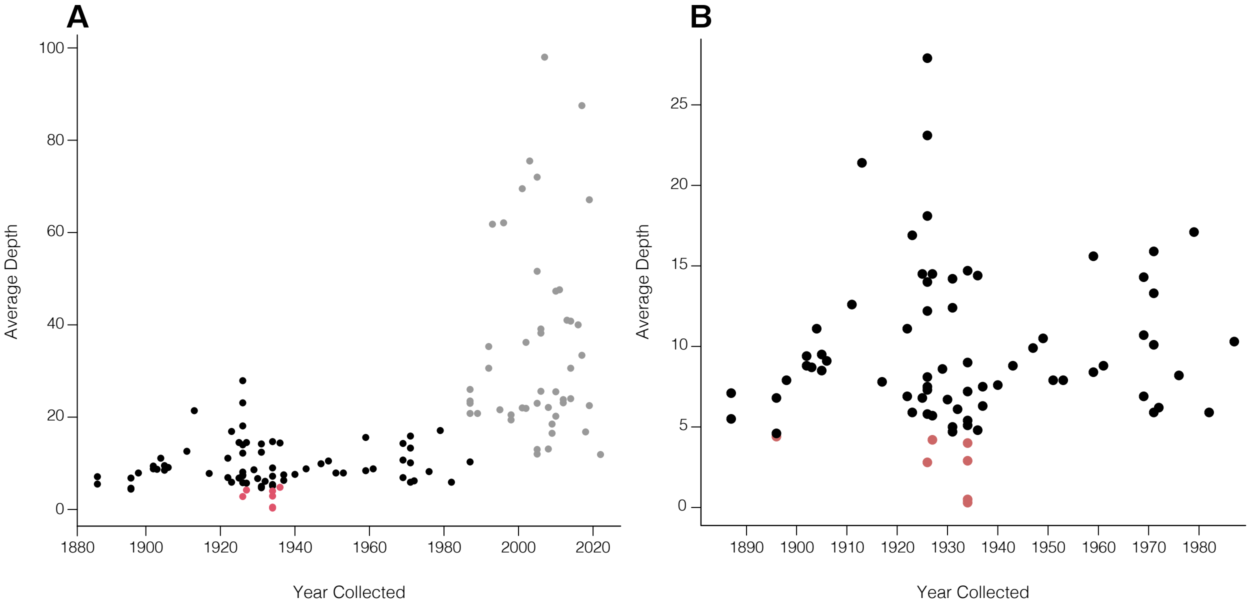


Fig S5. Average depth of all samples (A) and just toepad-sourced samples (B). Dot color indicate source of sample; gray dots are tissue-sourced, black dots are toepad-sourced, and red dots are toepad-sourced samples that were sequenced and successfully assembled but were not included in downstream analsyes.

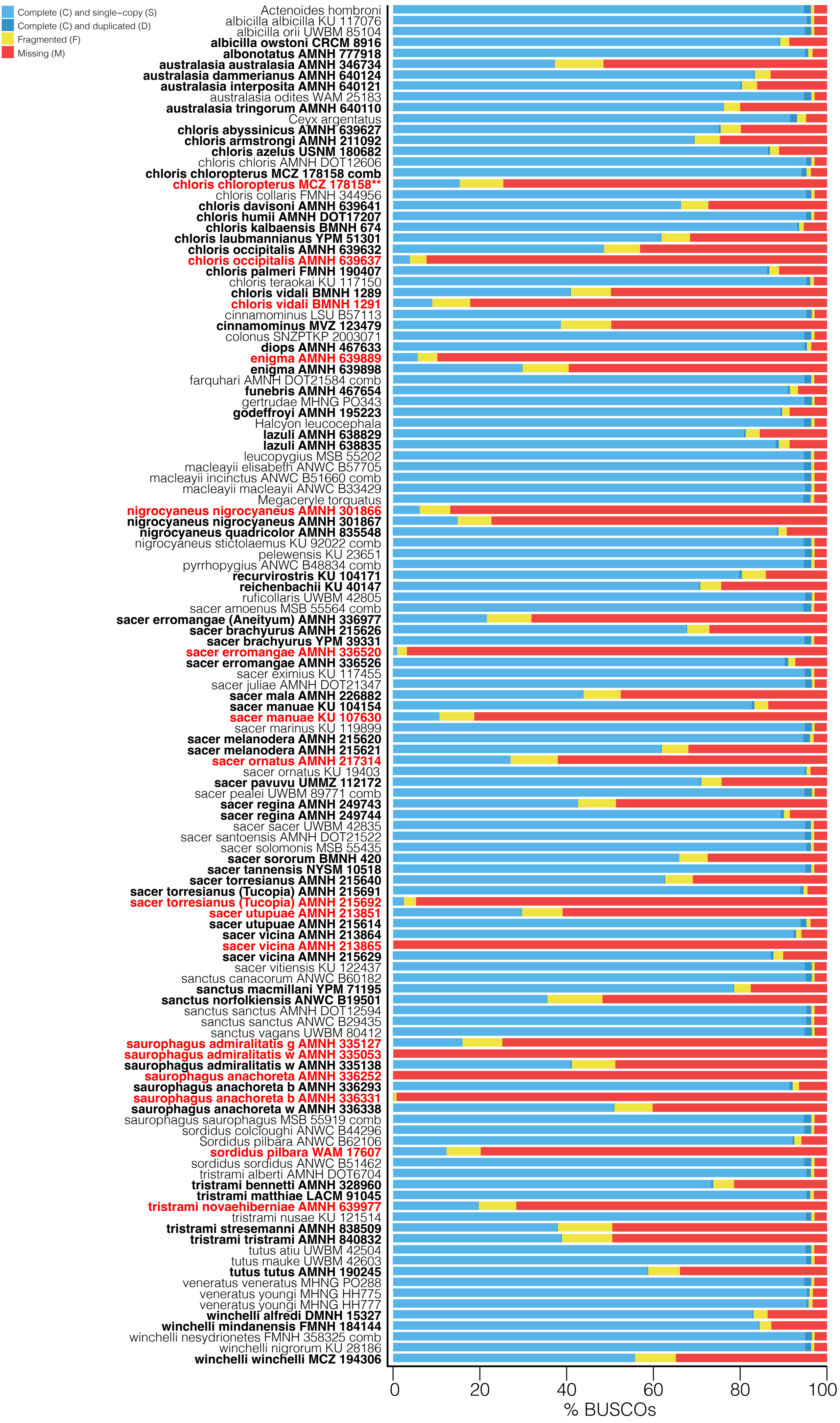


Figure S6: BUSCO completeness graph of all samples in this study. Light blue indicates complete and single-copy loci. Dark blue is complete and duplicated loci. Yellow is fragmented loci. Red is missing loci. Sample names that are bolded are toepad-sourced samples and red are samples that were not included in the BUSCO dataset.

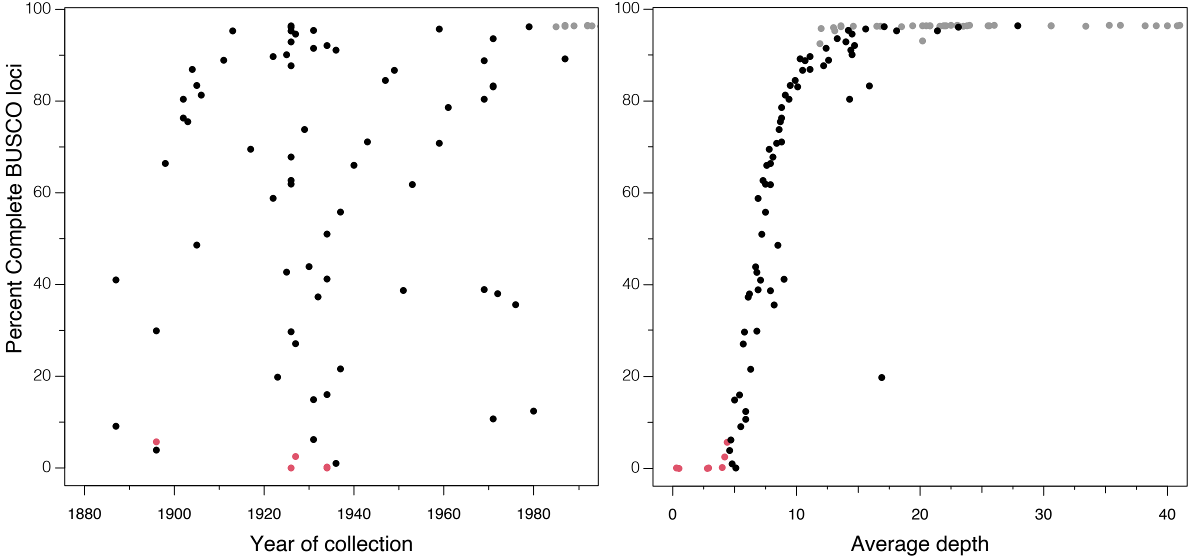


Figure S7: Average depth, not collection year, was more important for percent complete single-copy BUSCO loci for toepad-sourced genomes (out of 8,338 single-copy BUSCO loci). Dot color represents tissue-sourced (grey), toepad-sourced (red), and toepad-sourced samples that were not included in any subsequent analyses (red). We show only a subset of years and depths relevant to toepad-sourced genomes.

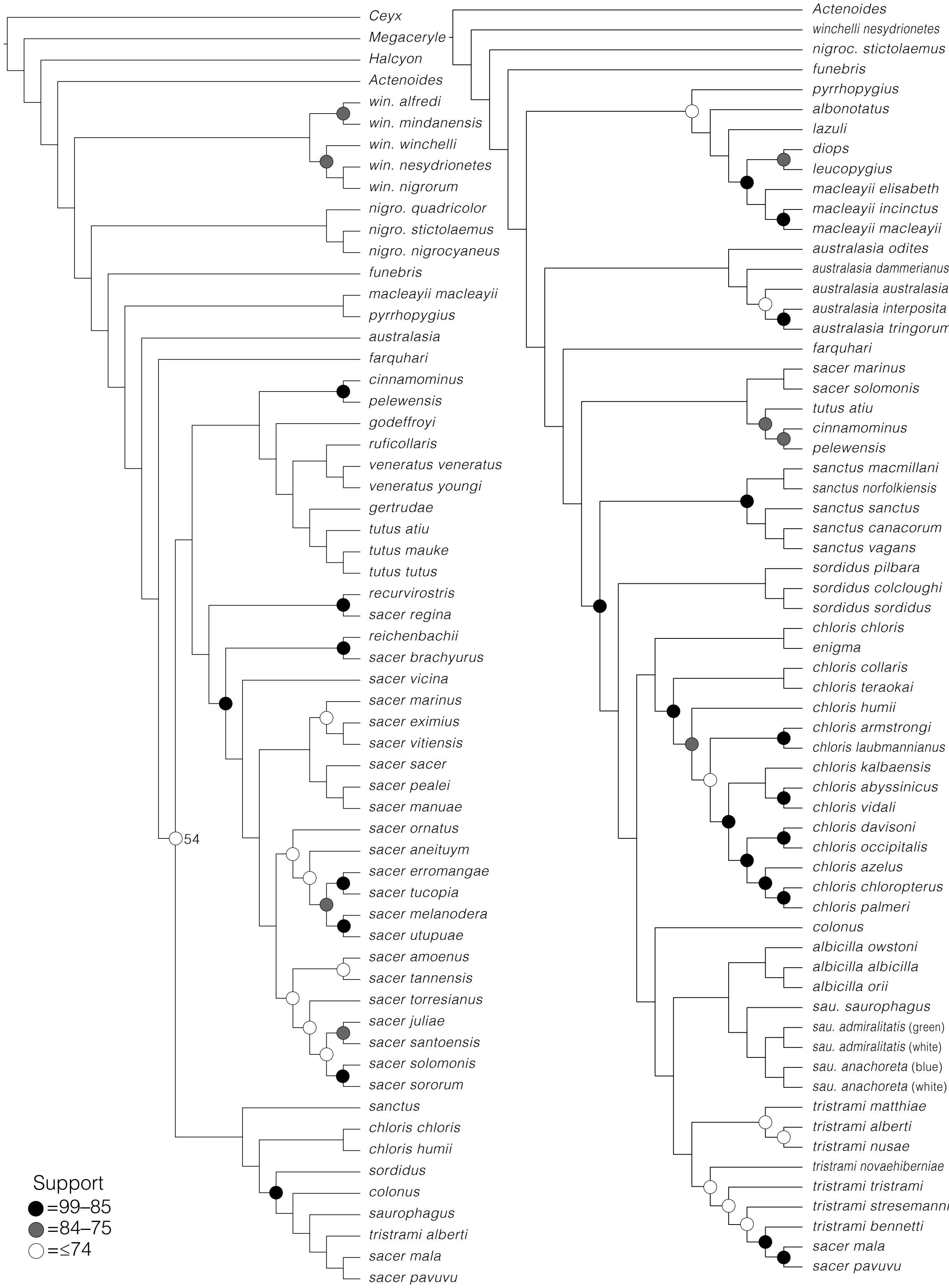


Figure S8: Subsets of the species tree, produced in SVDquartets, from the 90% complete UCE matrix (4,942 loci, 4,531,264 bp). Circles reflect node support values that are below 100 BS: black circles indicate 99–85 BS, gray circles indicate 84–75 support, and white circles indicate 74 and lower support.

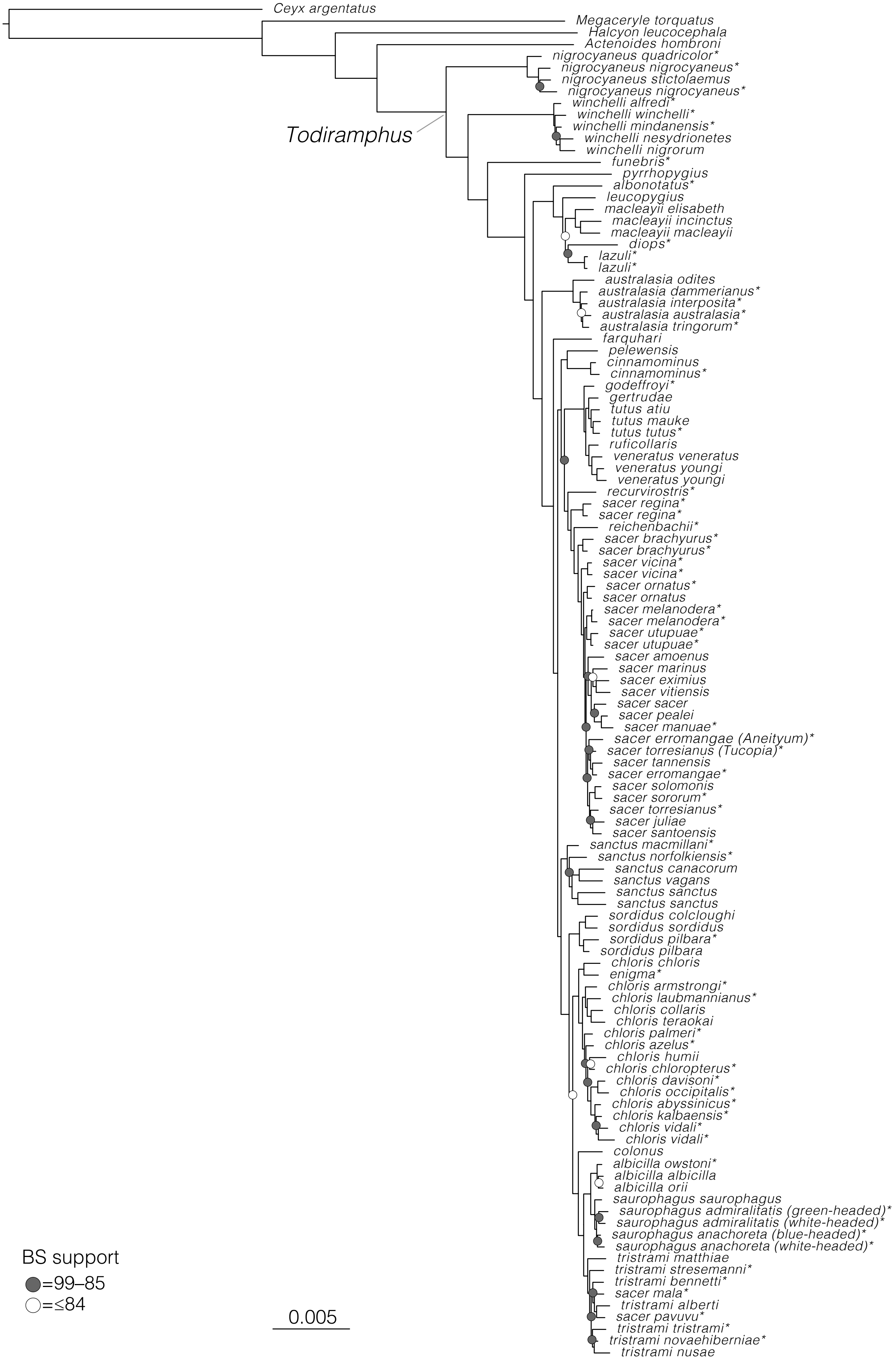


Figure S9: Partitioned maximum likelihood phylogeny, produced in IQtree, from the 90% complete UCE matrix (4,942 loci, 4,531,264 bp). Circles reflect node support values that are below 100 BS: gray circles indicate 99–85 support and white circles indicate 84 or lower support. Samples with asterisks (*) indicate toepad-sourced samples.

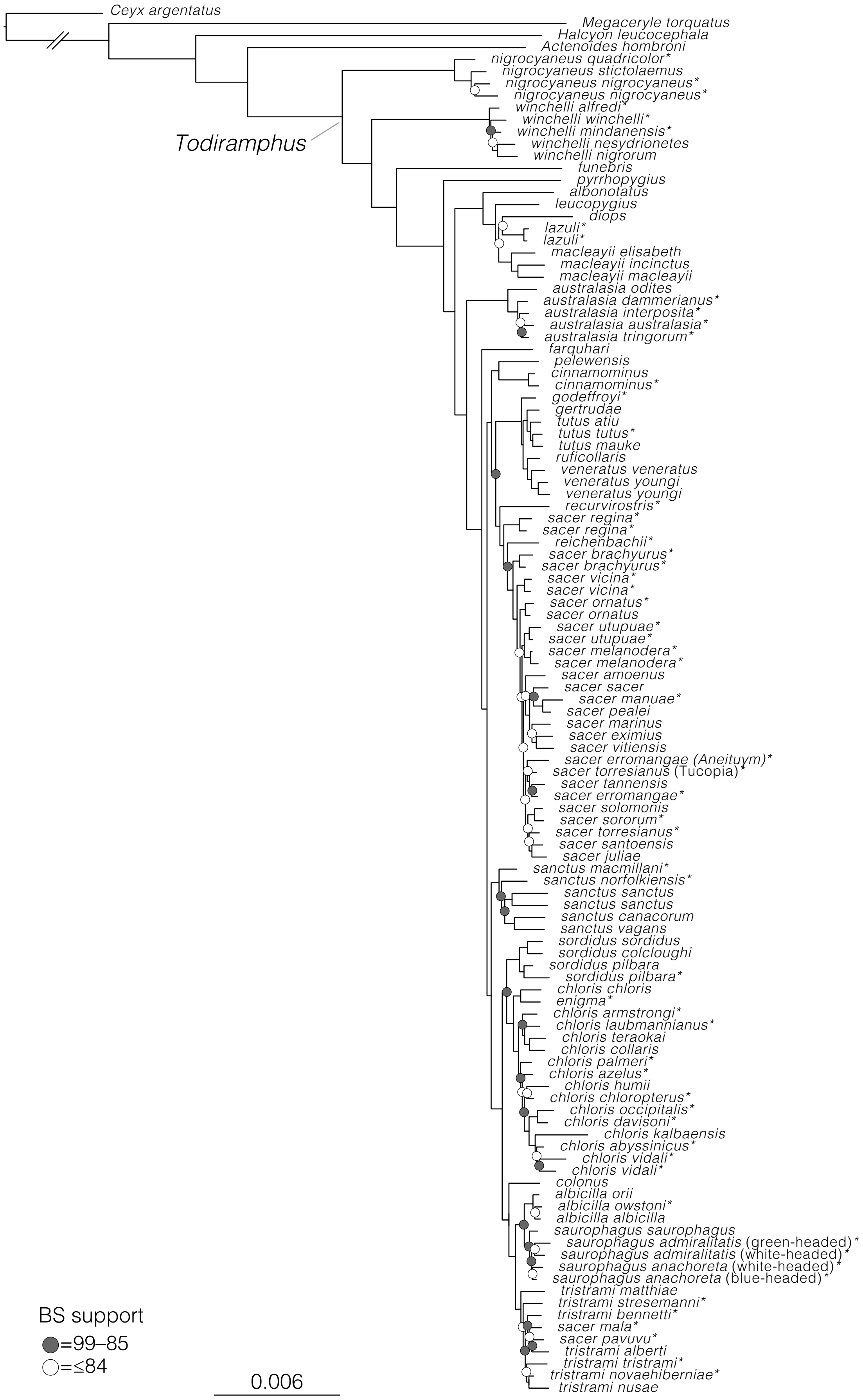


Figure S10: Maximum likelihood phylogeny, produced in RAxML-ng, from the 90% complete UCE matrix (4,942 loci, 4,531,264 bp). Circles reflect node support values that are below 100 BS: gray circles indicate 99–85 support and white circles indicate 84 or lower support.

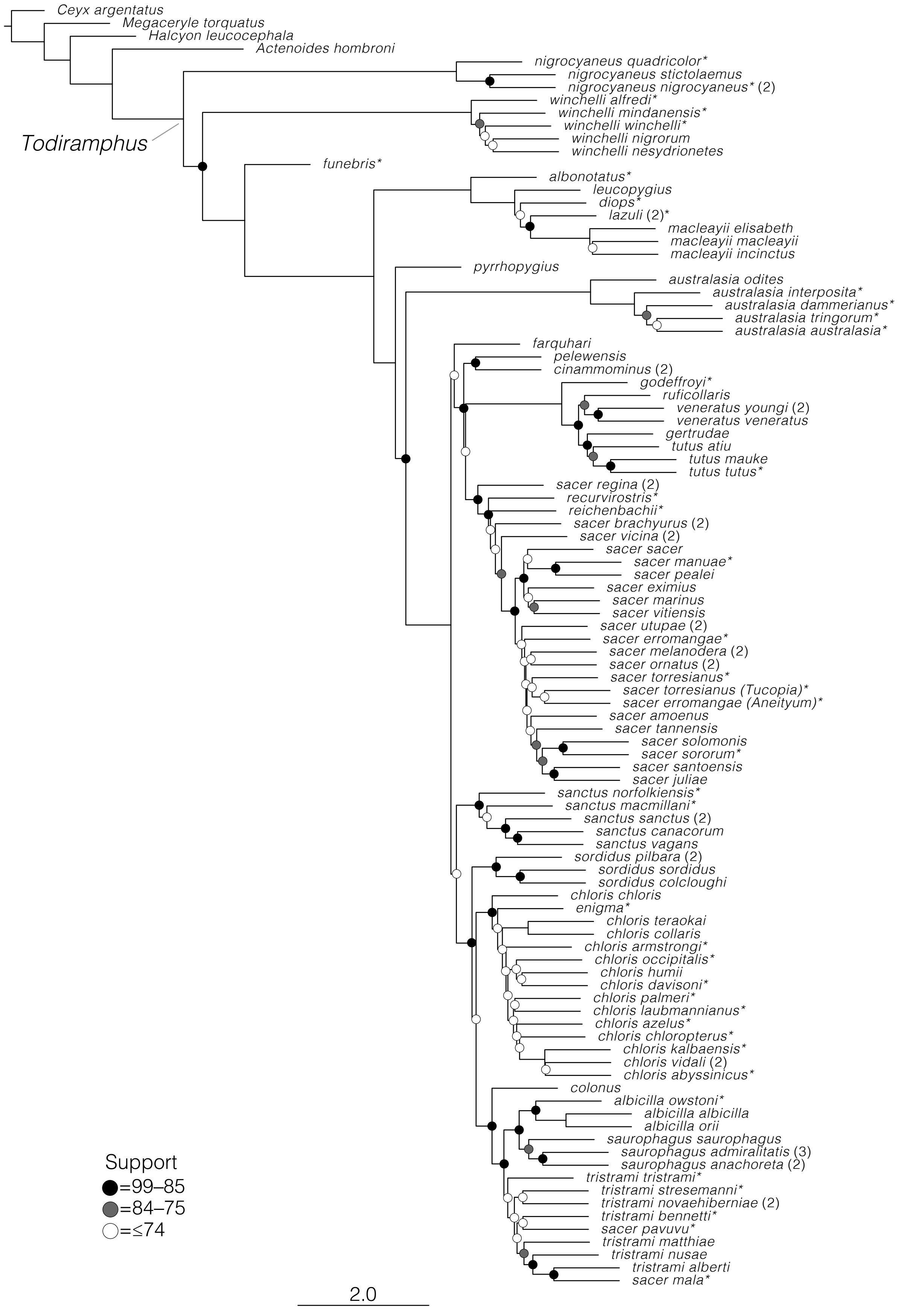


Figure S11: Species tree of Todiramphus kingfishers, produced in ASTRAL-III, based off of loci from the 90% complete that had at least 100 parsimony informative sites (125 loci). Circles reflect node support values that are below 100 BS: black circles indi- cate 99–85 BS, gray circles indicate 84–75 support, and white circles indicate 74 and lower support. Samples with asterisks (*) indicate toepad-sourced samples. Tips with numbers in parentheses indicate multiple samples that were distinct in gene trees but were lumped as a single tip for ASTRAL.

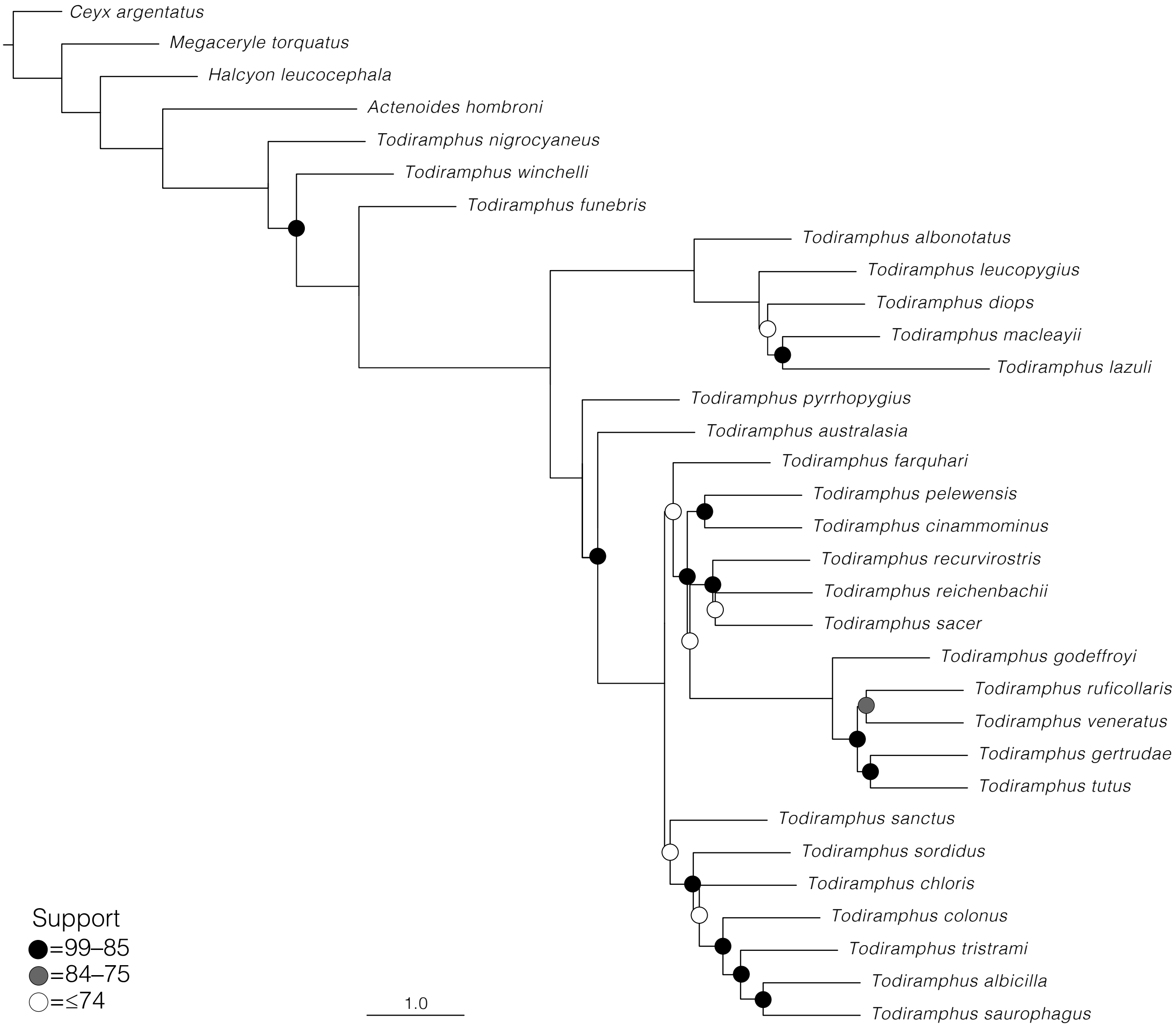


Figure S12: Species tree of Todiramphus kingfishers, produced in ASTRAL-III, based off of loci from the 90% complete that had at least 100 parsimony informative sites (125 loci). Circles reflect node support values that are below 100 BS: black circles indi- cate 99–85 BS, gray circles indicate 84–75 support, and white circles indicate 74 and lower support. See Fig S11 for all subspecific diversity.

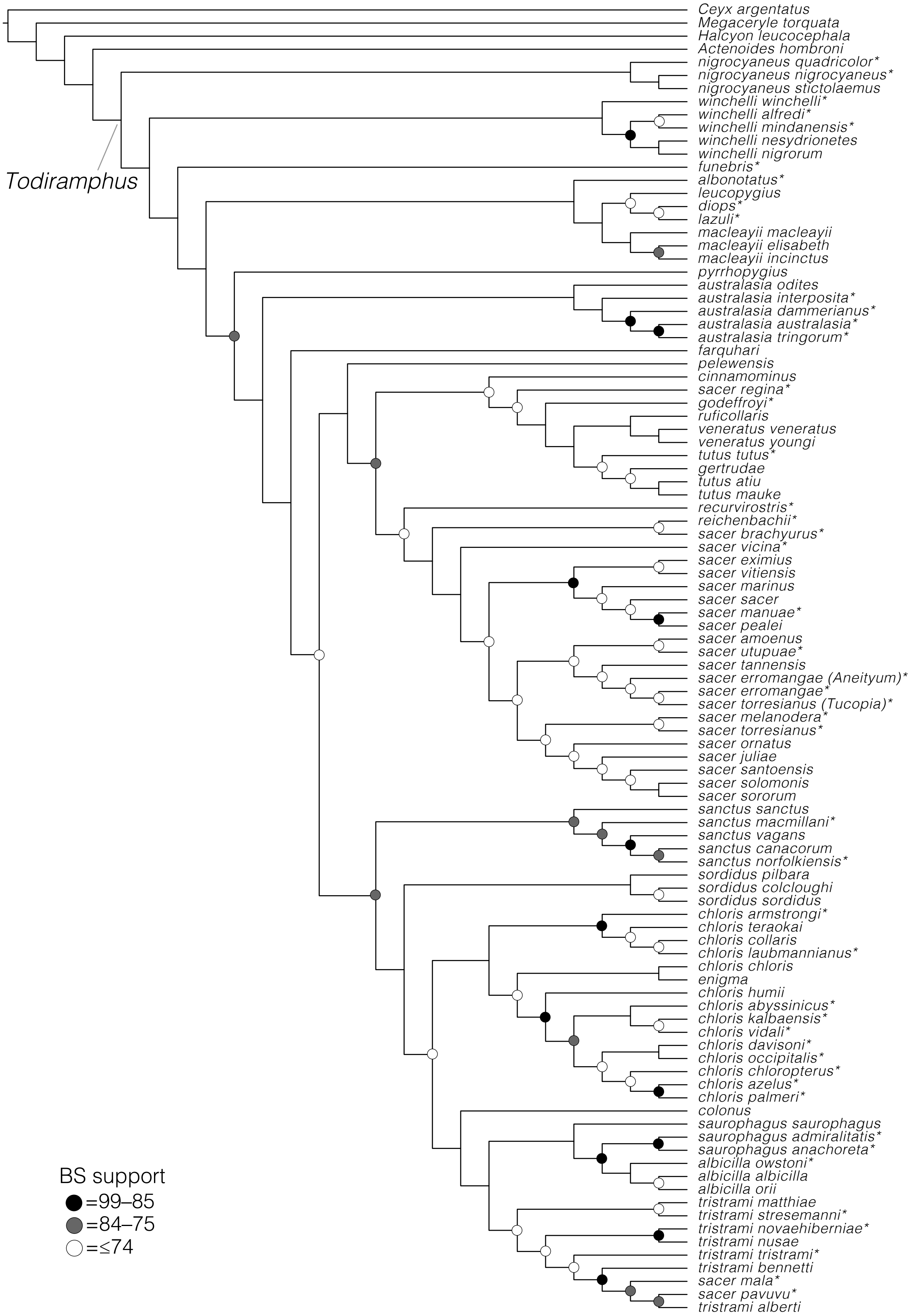


Figure S13: SNP-based species tree, produced in SVDquartets, from the 100% complete SNP dataset (85,156 SNPs) for 119 tips. Circles reflect node support values that are below 100 BS: black circles indicate 99–85 BS, gray circles indicate 84–75 support, and white circles indicate 74 and lower support. Samples with asterisks (*) indicate toepad-sourced samples.

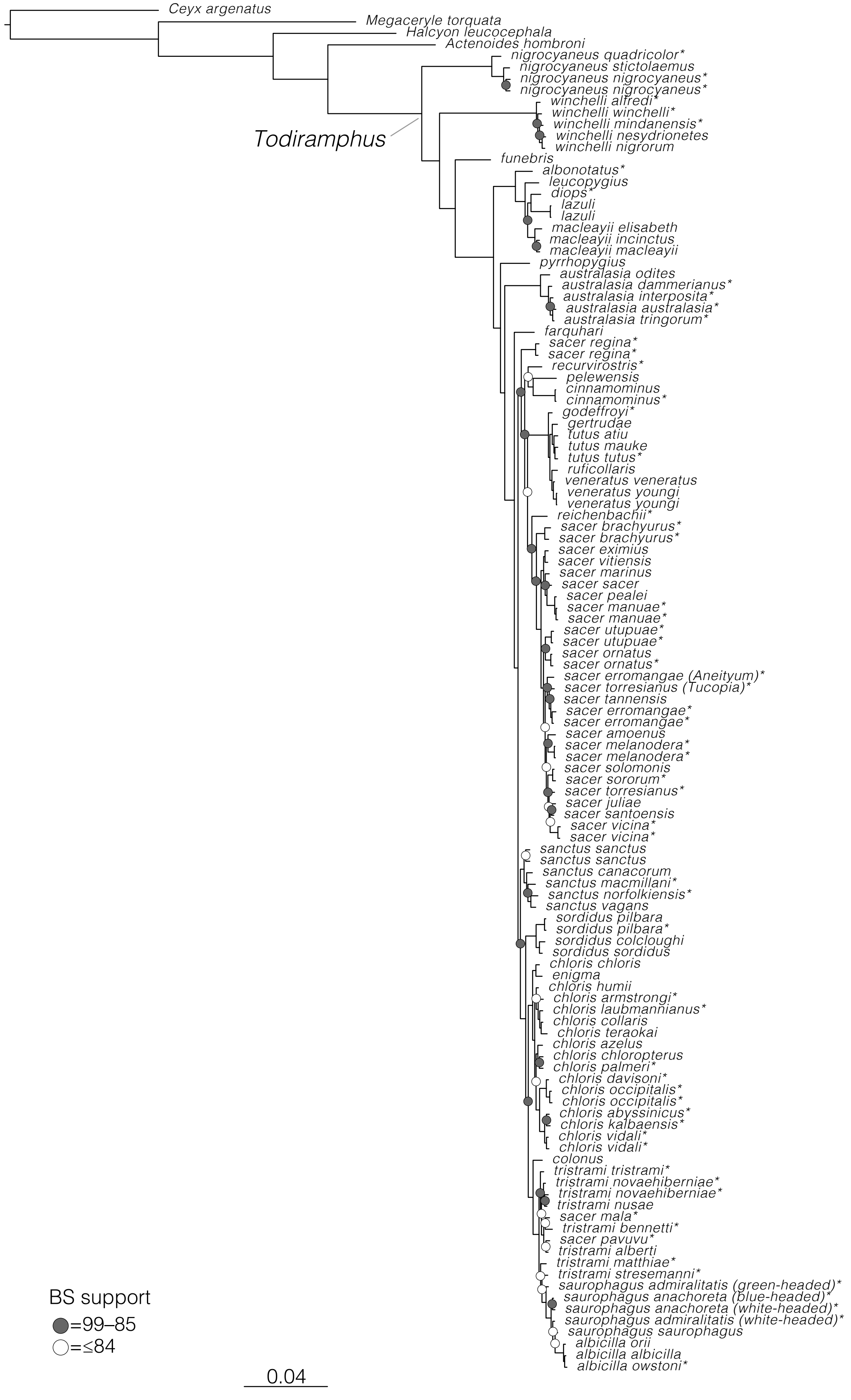


Figure S14: Maximum likelihood phylogeny, produced in IQtree, from the 100% complete SNP dataset (85,156 SNPs) for 119 tips. Circles reflect node support values that are below 100 BS: gray circles indicate 99–85 support and white circles indicate 84 or lower support. Samples with asterisks (*) indicate toepad-sourced samples.

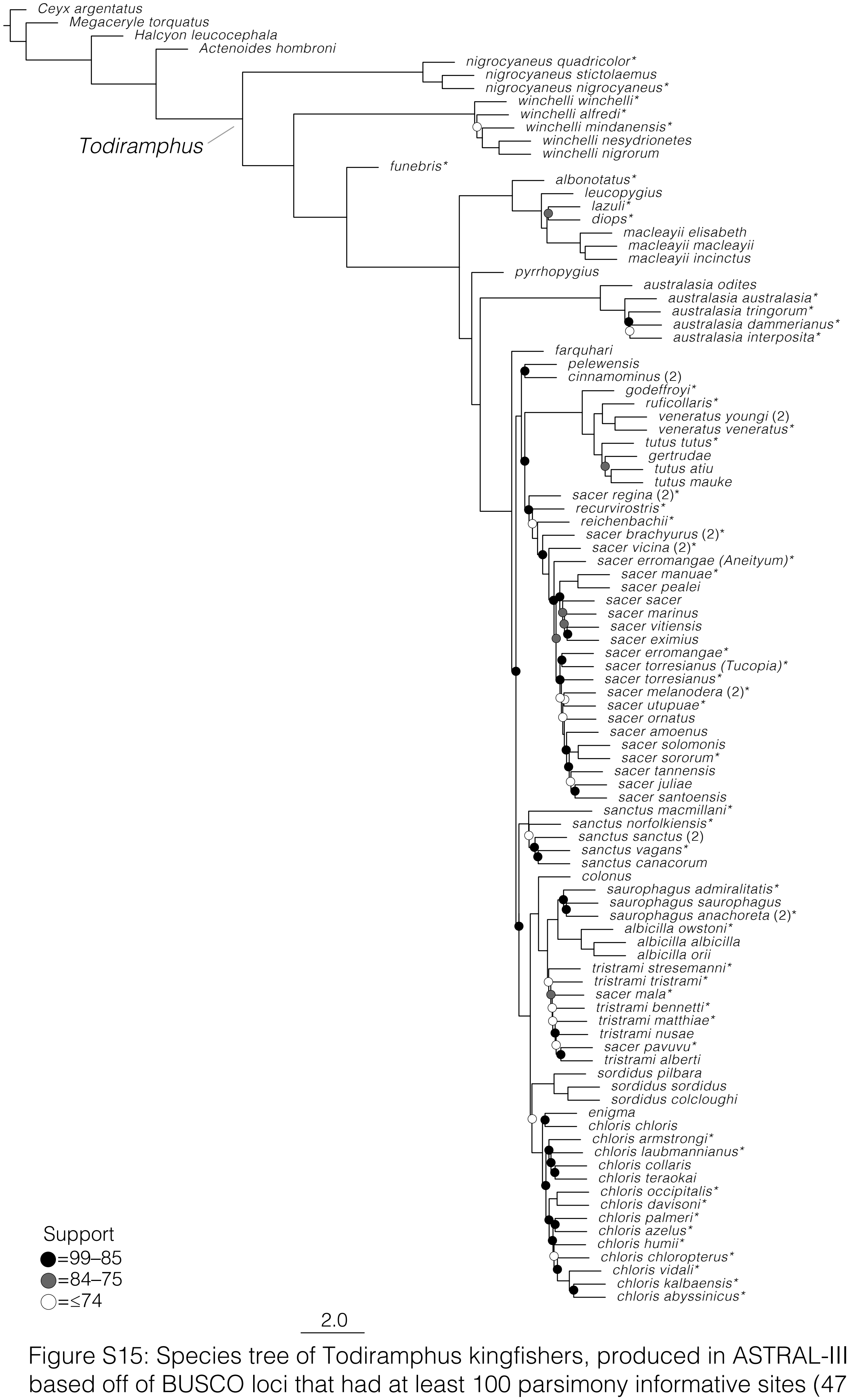


Figure S15: Species tree of Todiramphus kingfishers, produced in ASTRAL-III, based off of BUSCO loci that had at least 100 parsimony informative sites (470 loci, 815,282 AA sites). Circles reflect node support values that are below 100 BS: gray circles indicate 99–85 support and white circles indicate 84 or lower support. Samples with asterisks (*) indicate toepad-sourced samples.

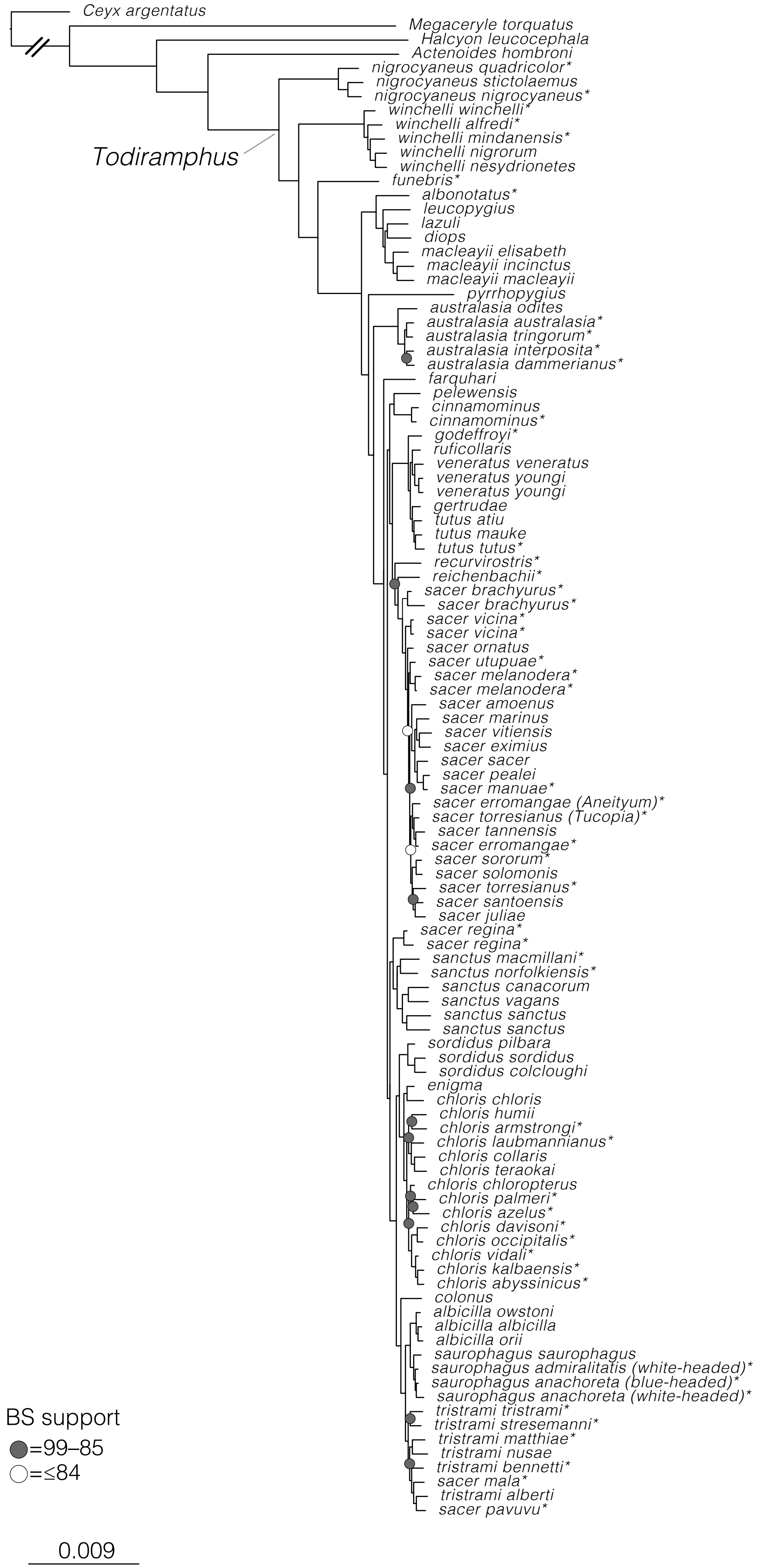


Figure S16: Partitioned maximum likelihood phylogeny, produced in IQtree, from the BUSCO dataset (8,012 loci, 5,102,241 AA sites). Circles reflect node support values that are below 100 BS: gray circles indicate 99–85 support and white circles indicate 84 or lower support. Samples with asterisks (*) indicate toepad-sourced samples.

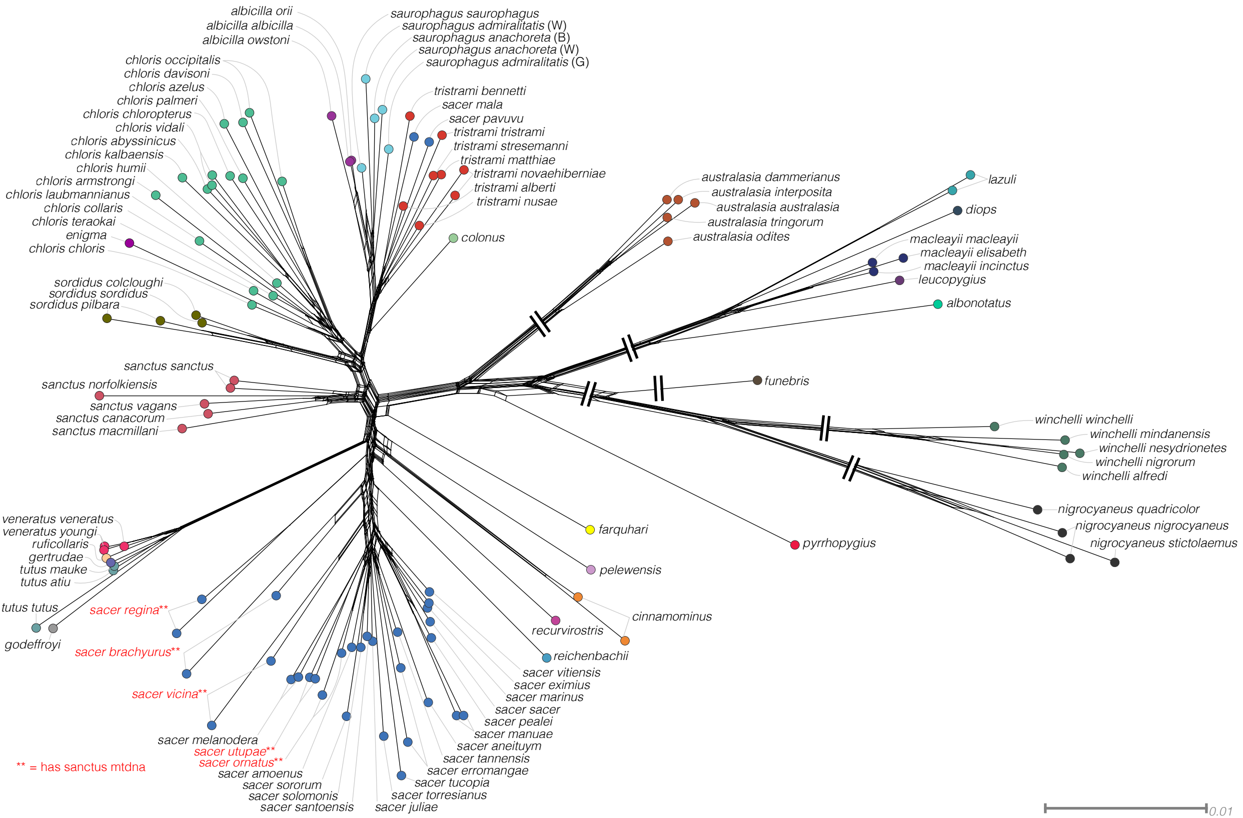


Figure S17: Unrooted phylogenetic network based on pairwise divergence among all samples, produced from a complete set of 85,165 SNPs. Note that some branch lengths have been artificially shortened, denoted by double slashes.

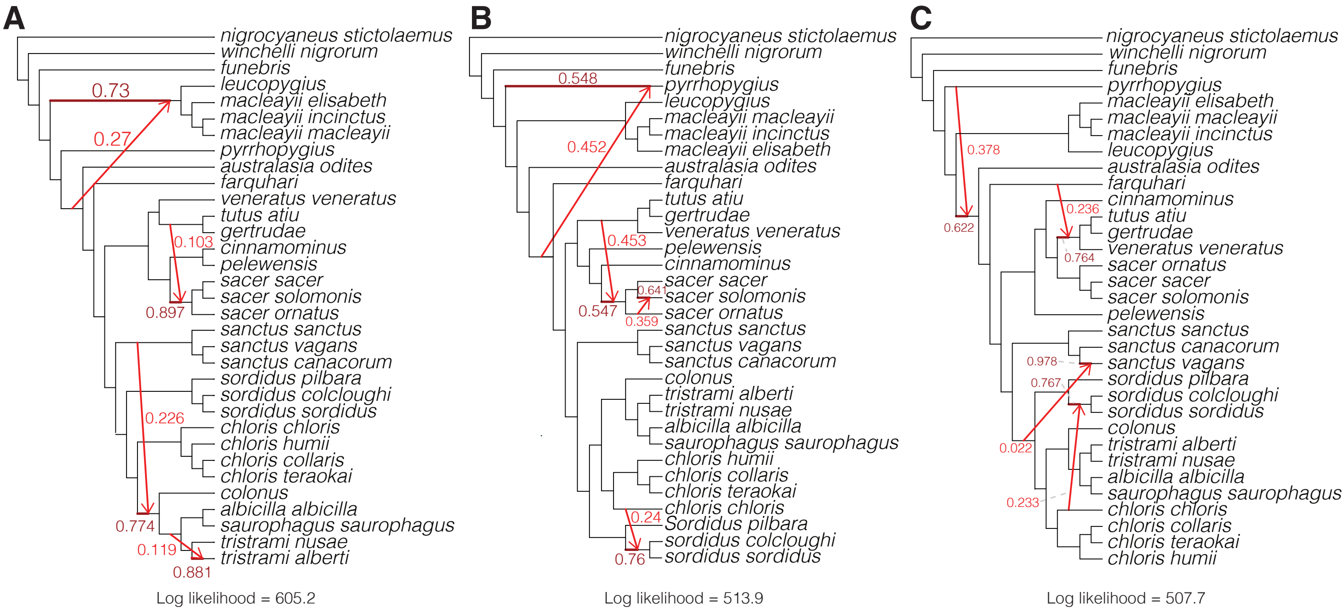


Figure S18: Phylogenetic networks, estimated in SNaQ, from three unique quartet subsets (3,000 independent quartets each). Red branches and arrows indicate the proportion of genes are inherited at reticulation events. Lighter red indicates inheritance values γ.

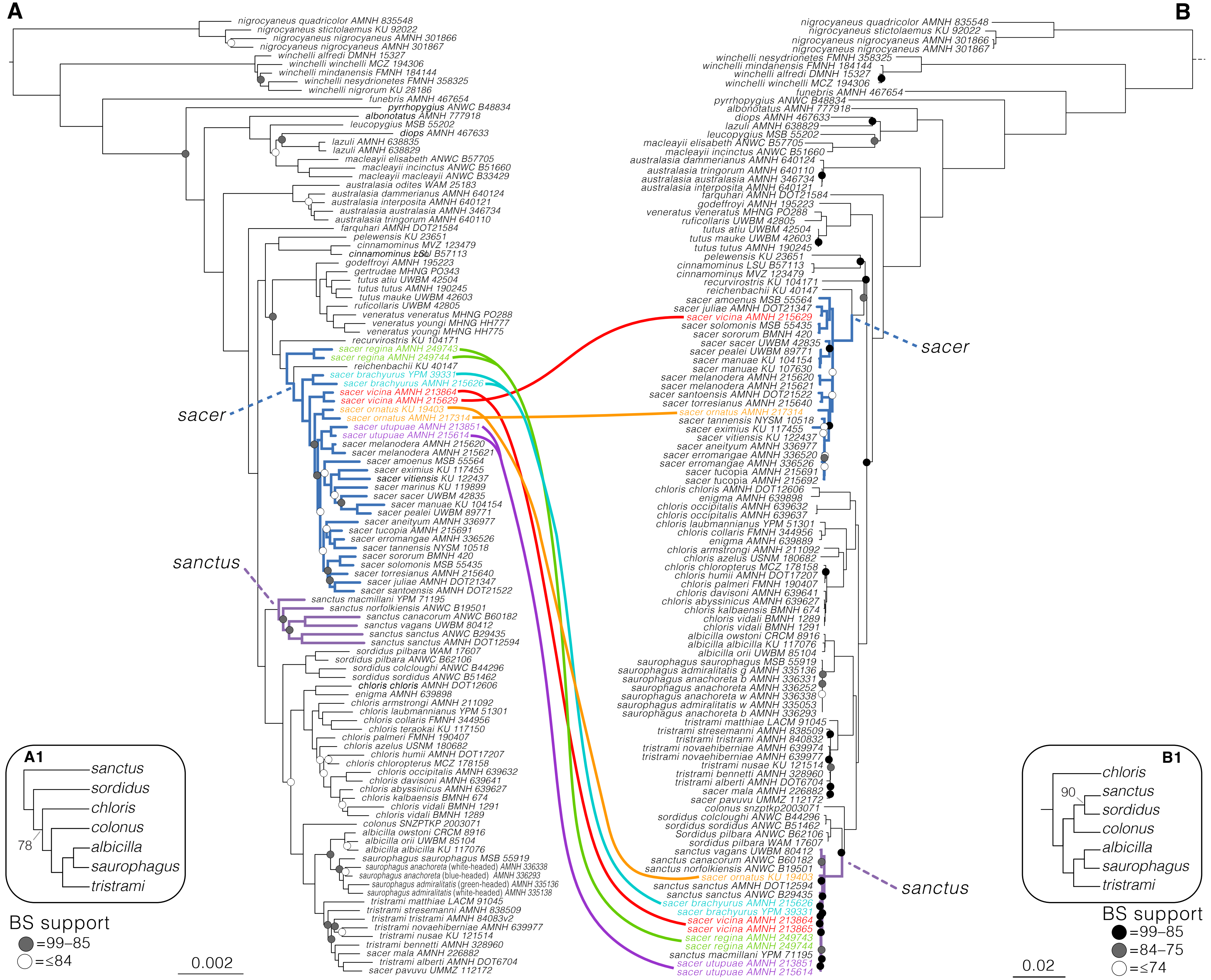


Figure S19. Mitochondrial capture within some Pacific Kingfisher (*T. sacer*) taxa and Sacred Kingfisher (*T. sanctus*). A) Partitioned IQtree topology inferred from the 90% complete UCE matrix. B) Partitioned maximum likelihood phylogeny inferred from 13 mitochondrial genes with IQtree. Note that the mtDNA phylogeny (B1) for the Australasian clade is different than the UCE topology (A1), which has been described previously (DeRaad et al. 2023). All cases *T. sacer* taxa with *T. sanctus* mtDNA are those that are endemic to small islands in Temotu Province, Solomon Islands (*T. s. ornatus, brachyurus, vicina, utupuae*) or Wallis and Futuna territory (*T. s. regina*). See Figs. 4 and S1 for taxon-specific maps.

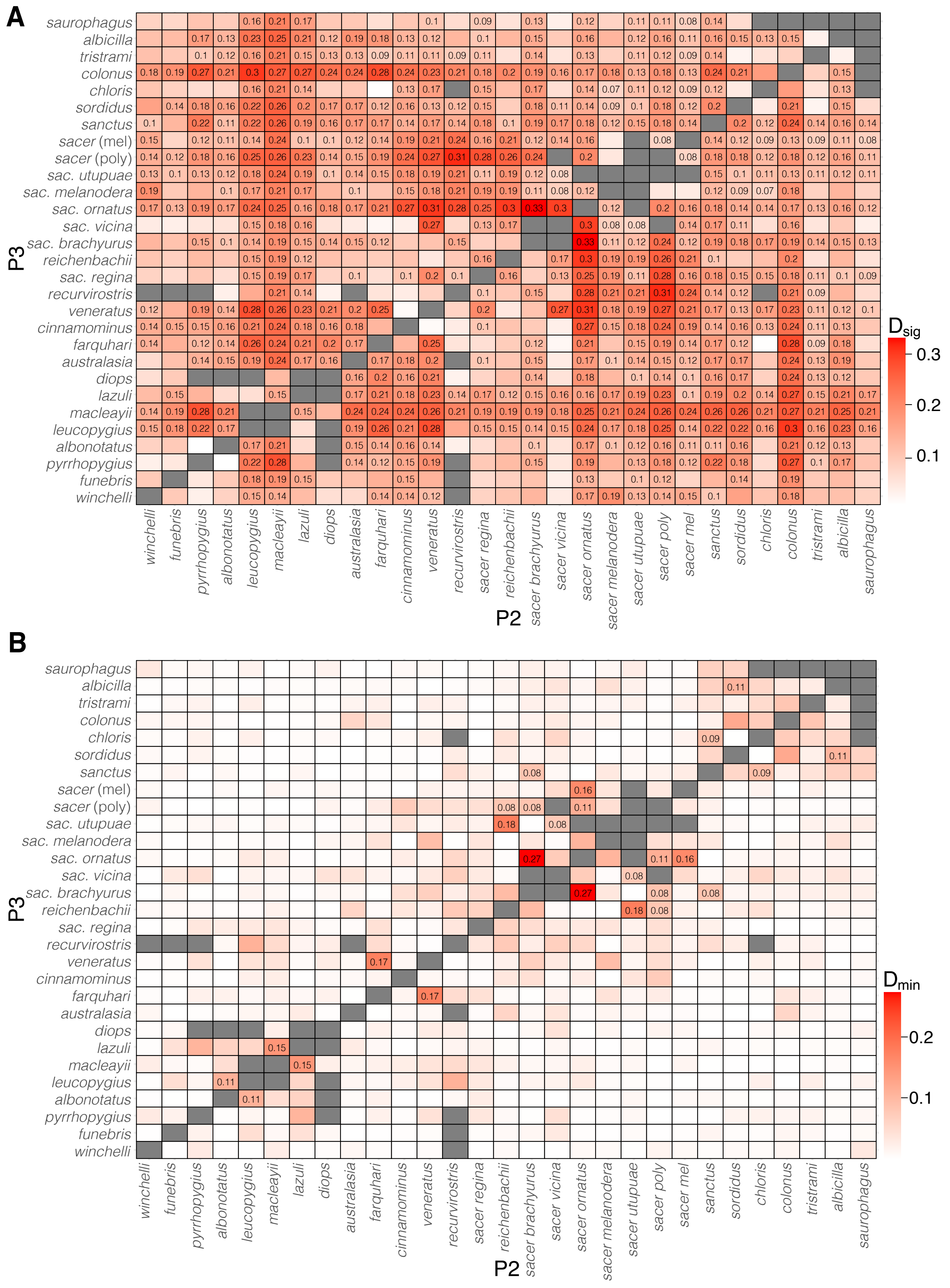


Figure S20: Pairwise results of ABBA-BABA tests, with text indicating D-statistics only in squares with FDR-adjusted significant pairwise p values. Panel A is the most significant (lowest pvalue) for any pairwise comparison and Panel B is the lowest value of D-statistics (Dmin). Here, *T. sacer* taxa that have *T. sanctus* mtDNA are considered their own tips and two *T. sacer* clades are those recovered in Figs. 1, S9.

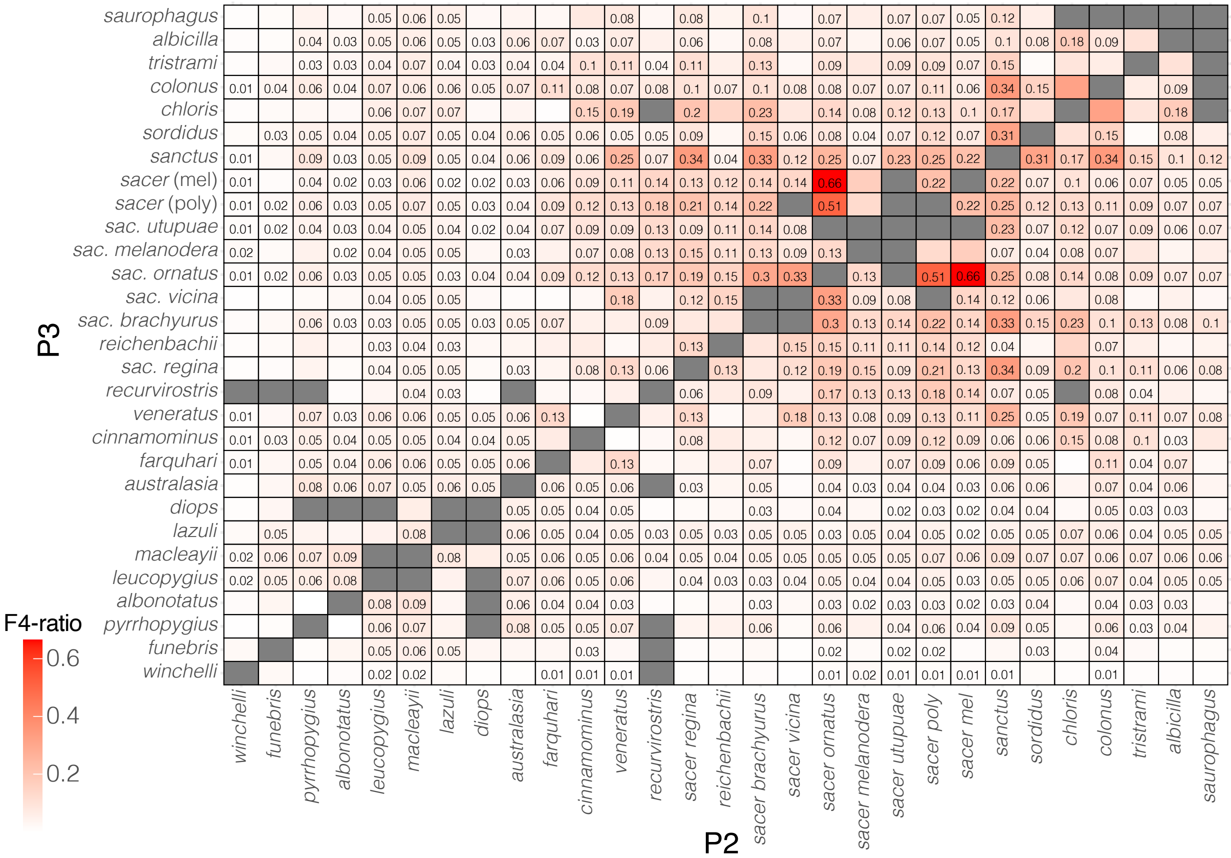


Figure S21: Pairwise results of ABBA-BABA tests, with text indicating F4-ratio values only in squares with FDR-adjusted significant pairwise p values (<0.000013). Here, *T. sacer* taxa that have *T. sanctus* mtDNA are considered their own tips and two *T. sacer* clades are those recovered in Fig. 1.

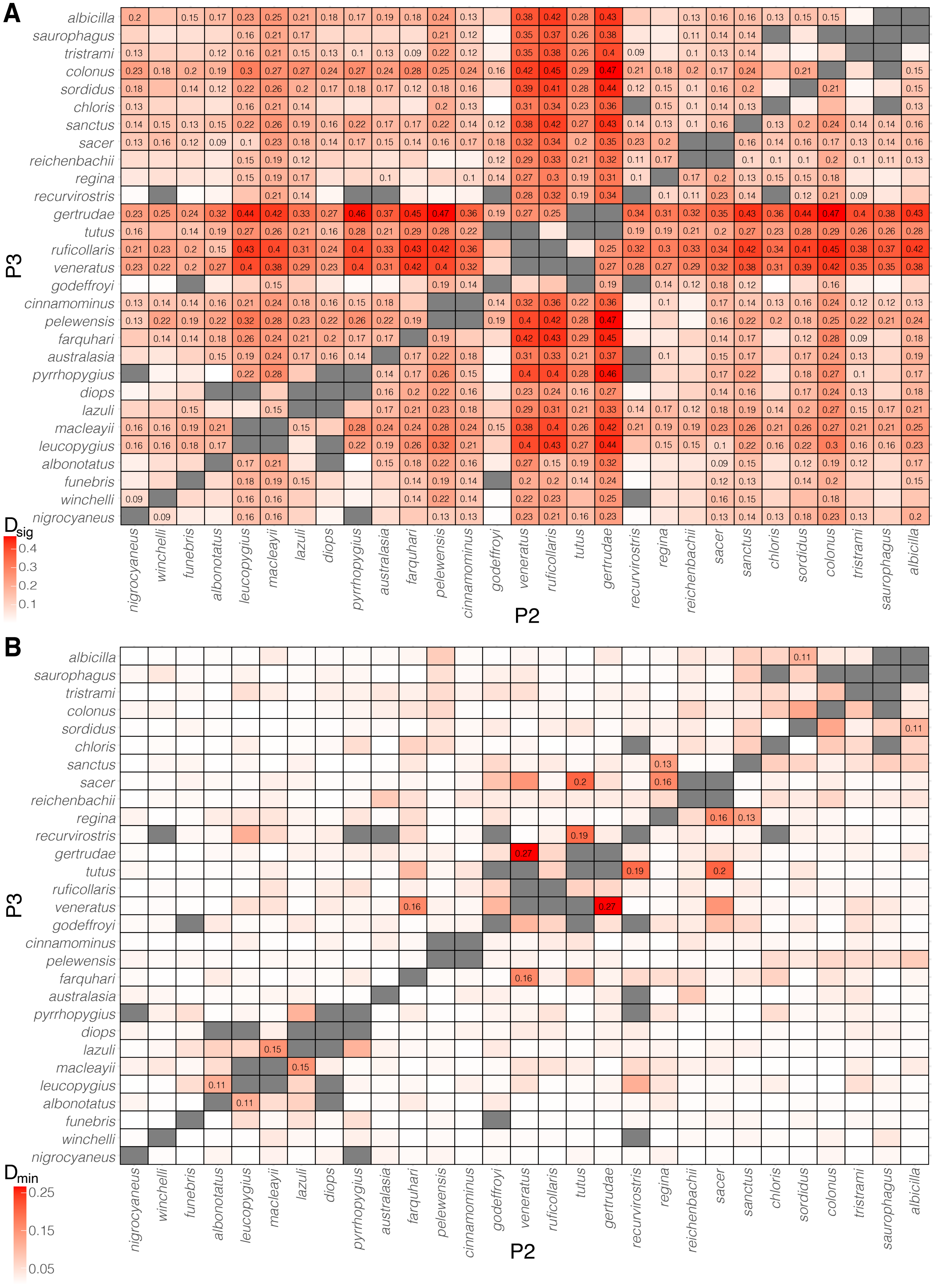


Figure S22: Pairwise results of ABBA-BABA tests, with text indicating D-statistics only in squares with FDR-adjusted significant pairwise p values (<0.000013). Panel A is the most significant (lowest pvalue) for any pairwise comparison and Panel B is the lowest value of D-statistics (Dmin). Here, only *T. s. regina* is considered its own tip and all other T. sacer taxa are lumped into a simple tip.

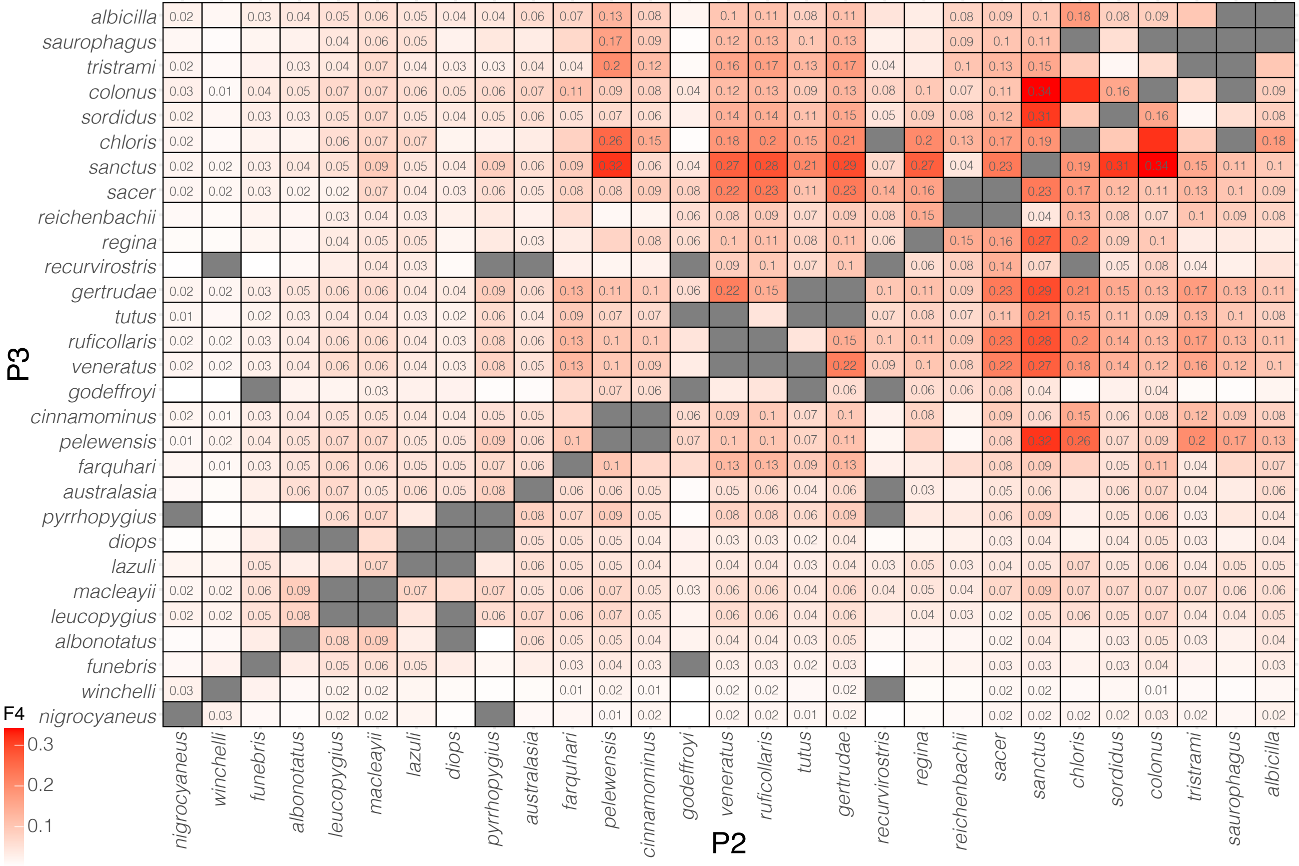


Figure S23: Pairwise results of ABBA-BABA tests, with text indicating F4-ratio values only in squares with FDR-adjusted significant pairwise p values (<0.000013). Here, only *T. s. regina* is considered its own tip and all other *T. sacer* taxa are lumped into a simple tip.
